# Distinct hydrogenotrophic bacteria are stimulated by elevated H_2_ levels in upland and wetland soils

**DOI:** 10.1101/2020.11.15.383943

**Authors:** Yongfeng Xu, Ying Teng, Xiyang Dong, Xiaomi Wang, Chuwen Zhang, Wenjie Ren, Ling Zhao, Yongming Luo, Chris Greening

## Abstract

**Background:** Molecular hydrogen (H_2_) is a major energy source supporting bacterial growth and persistence in soil ecosystems. While recent studies have uncovered mediators of atmospheric H_2_ consumption, far less is understood about how soil microbial communities respond to elevated H_2_ levels produced through natural or anthropogenic processes. Here we performed microcosm experiments to resolve how microbial community composition, capabilities, and activities change in upland (meadow, fluvo-aquic soil) and wetland (rice paddy, anthrosols soil) soils following H_2_ supplementation (at mixing doses from 0.5 to 50,000 ppmv).

**Results:** Genome-resolved metagenomic profiling revealed that these soils harbored diverse bacteria capable of using H_2_ as an electron donor for aerobic respiration (46 of the 196 MAGs from eight phyla) and carbon fixation (15 MAGs from three phyla). H_2_ stimulated the growth of several of these putative hydrogenotrophs in a dose-dependent manner, though the lineages stimulated differed between the soils; whereas actinobacterial lineages encoding group 2a [NiFe]-hydrogenases grew most in the upland soils (i.e. Mycobacteriaceae, Pseudonocardiaceae), proteobacterial lineages harboring group 1d [NiFe]-hydrogenases were most enriched in wetland soils (i.e. Burkholderiaceae). Hydrogen supplementation also influenced the abundance of various other genes associated with biogeochemical cycling and bioremediation pathways to varying extents between soils. Reflecting this, we observed an enrichment of a hydrogenotrophic *Noviherbaspirillum* MAG capable of biphenyl hydroxylation in the wetland soils and verified that H_2_ supplementation enhanced polychlorinated biphenyl (PCB) degradation in these soils, but not the upland soils.

**Conclusions:** Our findings suggest that soils harbour different hydrogenotrophic bacteria that rapidly grow following H_2_ exposure. In turn, this adds to growing evidence of a large and robust soil H_2_ sink capable of counteracting growing anthropogenic emissions.

## Background

Recent work has revealed that molecular hydrogen (H_2_) oxidation is a widespread treat among soil bacteria [1–4]. H_2_ is ubiquitously available in all soils through atmospheric and edaphic sources [5, 6]. Bacteria expend few resources to mobilize this gas, given both its diffusivity through cell membranes and low activation energy, and can use the large amount of free energy released by its oxidation for both ATP synthesis and carbon dioxide (CO_2_) fixation [2, 7, 8]. Genomic and metagenomic surveys have shown that soil bacteria from at least 17 different phyla encode [NiFe]-hydrogenases to consume H_2_ an energy source [1–3, 9, 10]. The most abundant H_2_-oxidising taxa in oxygenated soils are generally Actinobacteriota, Acidobacteriota, and Chloroflexota that encode high-affinity group 1h [NiFe]-hydrogenases [11–19]; culture-based studies show these bacteria use this enzyme to scavenge trace concentrations of H_2_ as an alternative energy source for persistence when organic growth substrates are limiting [12, 15–17, 20–24]. A smaller proportion of soil bacteria can grow autotrophically or mixotrophically on H_2_/CO_2_ [1, 3, 25–27]. Classical studies have shown numerous Proteobacteria, for example *Ralstonia eutropha* and *Bradyrhizobium japonicum*, grow efficiently on high levels of H_2_ using the low-affinity group 1d [NiFe]-hydrogenases [27–34]. More recently, diverse taxa have been shown to use group 2a [NiFe]-hydrogenases to grow on H_2_ at a wide range of concentrations [35–39]. Some bacteria use H_2_ for multiple purposes; for example, some *Mycobacterium* species switch between synthesising the growth-supporting 2a hydrogenase and persistence-supporting 1h hydrogenase in response to organic carbon availability [40–42].

Despite these advances, we lack a sophisticated understanding of how soil microbial communities respond to H_2_ availability. In most soils, bacteria are primarily exposed to H_2_ at atmospheric mixing ratios (~0.53 ppmv) [43, 44]. Soil bacteria use high- and medium-affinity hydrogenases to consume this trace energy source during growth or survival [3, 17, 37, 45–47]. Through their activity, approximately 70 million tonnes of the net H_2_ lost from the atmosphere each year, with far-reaching ecological and biogeochemical consequences [6, 48–50]. This process supports the productivity and diversity of bacteria, especially in oligotrophic environments [3, 18, 51, 52]. Moreover, it serves as the main sink in the global hydrogen cycle, in turn regulating the redox state and greenhouse gas levels of the atmosphere [6, 50, 53]. Nevertheless, multiple environments are known where H_2_ availability is elevated, for example due to biological fermentation and nitrogen fixation, or geological processes. For example, H_2_ can accumulate to percentage levels (~20,000 ppmv) at the interface of soils and root nodules, as a result of obligate H_2_ production during the nitrogenase reaction [2, 54, 55]. In turn, these emissions have been proposed to influence rhizosphere microbial composition and potentially even fertilise plant growth [33, 56–59]. Furthermore, H_2_ emissions have also been proposed to enhance bioremediation of organochloride pollutants through direct or indirect mechanisms [60–62]. However, it has proven highly challenging to disentangle the effects of H_2_ exposure on microbial composition and activity in field settings from other variables. Similarly, unresolved is to what extent soil microbial communities can respond to anthropogenic H_2_ emissions [50]. It has been controversially proposed that the transition to a hydrogen economy would drastically increase atmospheric H_2_ levels and in turn induce climate forcing [63, 64]. Nevertheless, the microbial soil sink has so far maintained atmospheric H_2_ at constant levels, despite anthropogenic activities currently accounting for approximately half of net atmospheric H_2_ production [6]. Thus, it is essential to understand how soil microbial composition responds to elevated H_2_ to simultaneously resolve how this gas influences structure of natural ecosystems and predict responses to forecast emissions.

Several studies have used microcosms to investigate how soil microbial composition and activity changes following elevated H_2_ (eH_2_) exposure, albeit with strikingly different results. A shift in the biphasic kinetics of soil H_2_ uptake in response to elevated H_2_ [65–67]: the high-affinity H_2_ oxidation activities that dominate in untreated soils diminish in favour of fast-acting, low-affinity processes [67–70]. This suggests that low-affinity hydrogenotrophs become more abundant or active following H_2_ exposure, though there are apparent discrepancies as to which. A pioneering study by Osborne *et al* indicated that H_2_ production has a minimal effect on microbial abundance, composition, and diversity, but elicited a consistent enrichment of actinobacterial taxa across multiple soil types, including mycobacteria [57]. Zhang and colleagues, by contrast, observed Actinobacteriota decreased and Gammaproteobacteria increased following H_2_ exposure [71]. Given both of these restriction fragment length polymorphism (RFLP)-based studies predated current genome-resolved metagenomic approaches, the taxonomic identity and hydrogenase content of the enriched taxa could not be resolved. More recently, the Constant group have reexamined the effects of H_2_ supplementation using amplicon and metagenomic sequencing. They observed large-scaler differences in community composition and function between the treatment and control groups [67, 72]. They also reported that H_2_-oxidising taxa are rare community members and hence couldn’t be accurately accounted for even with deep metagenomic sequencing [73, 74]. Another recent study reported enrichment of ammonia-oxidising archaea and specific actinobacterial and acidobacterial lineages, as well as ammonia-oxidising archaea, following soil H_2_ infusion [75].

Altogether, these divergent observations warrant new investigations into the effects of H_2_ exposure on microbial community composition and activities. To do so, we investigated how H_2_ exposure at six different doses (from 0.5 to 50,000 ppmv) influences two agricultural soils from China with a legacy of organochloride pesticide usage, namely an anthrosols soil (herein wetland soil) and fluvo-aquic soil (herein upland soil). We combined high-resolution amplicon sequencing with deep genome-resolved metagenomic sequencing to resolve the taxonomic identities and metabolic capabilities of the taxa that change in abundance in response to H_2_ exposure. We show that both soils harbour a high abundance and diversity of H_2_-oxidising bacteria, and most taxa capable of autotrophic growth on H_2_/CO_2_ were generally enriched at higher H_2_ concentrations. However, reflecting differences in the community structure of the original soils, the enriched lineages strikingly differ in both phylogenetic affiliation and hydrogenase content between the upland and wetland microcosms. Contrasting changes in biogeochemical cycling genes and, building on our previous observations [60, 61], polychlorinated biphenyl (PCB) biodegradation processes were also observed between the soils following H_2_ exposure. Thus, the effects of H_2_ supplementation are highly ecosystem-specific, which reconciles the perplexingly different responses observed to H_2_ supplementation in studies in this area.

## Results and Discussion

### Elevated H_2_ stimulates growth of different bacteria between the soils, but does not significantly affect community richness or abundance

We first used the 16S rRNA gene as a marker to profile how abundance, alpha diversity, and beta diversity of bacteria and archaea present in the wetland and upland soils changed in response to H_2_ exposure. In agreement with the findings of Osborne et al [57], no significant change was observed in community abundance (based on 16S rRNA gene qPCR; **Fig. 1a & Fig. S1a**) or diversity (based on observed richness, Chao1 richness, and Shannon diversity of 16S rRNA gene amplicon sequence variants; **Fig. 1a & Fig. S2**) between the control and treatment microcosms. However, bacterial community composition changed in response to the H_2_ treatment after 84 days. Distance-based redundancy analysis (db-RDA) of beta diversity (Bray-Curtis of 16S rRNA gene amplicon sequence variants) confirmed H_2_ concentration is the predictor variable most significantly correlated with changes in bacterial community composition between the microcosms (*R*^2^ = 0.819, *p* = 0.001 in the wetland soil; *R*^2^ = 0.950, *p* = 0.001 in the upland soil; **Table S1**). For example, the samples treated with elevated H_2_ (500 to 50,000 ppmv in the wetland soil; 20,000 to 50,000 ppmv in the upland soil) formed distinct clusters from the control in PCoA data space (**Fig. 1b**). Three other predictor variables were also correlated with changes in community composition, most notably pH (*R*^2^ = 0.629, *p* = 0.001 in the wetland soil; *R*^2^ = 0.320, *p* = 0.047 in the upland soil; **Table S1**), which significant decreased during the H_2_-enriched microcosms likely as a result of soil bacteria oxidising H_2_ to protons **(Table 1)**.

**Figure 1.**
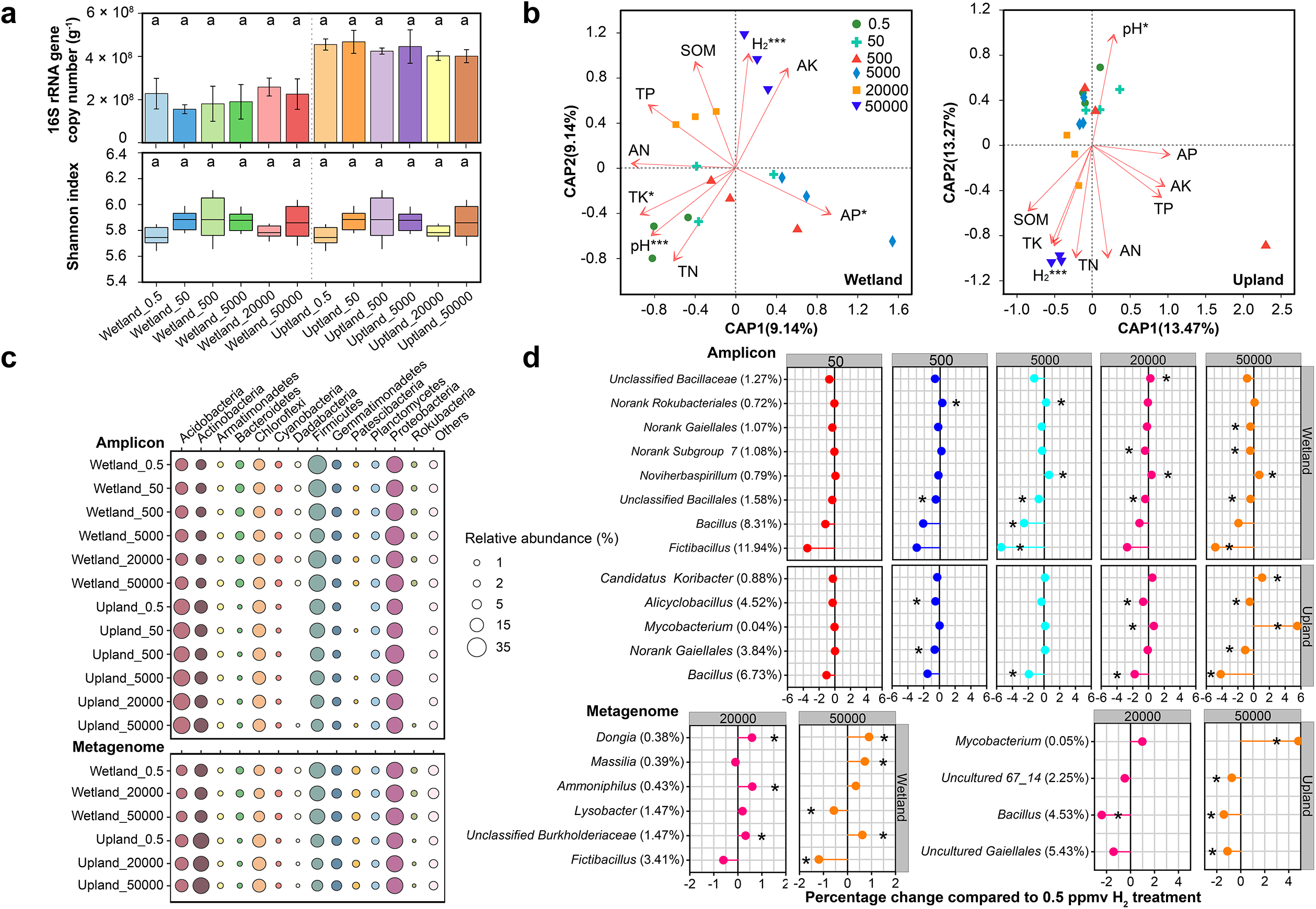
Changes of community abundance, diversity, and composition during the soil microcosms. **(a)** Stacked bar chart showing the estimated abundance of bacterial and archaeal taxa based on 16S rRNA gene copy number; Boxplot showing Shannon index of microbial communities based on 16S rRNA gene amplicon sequence variants. **(b)** The relationship between H_2_ mixing ratio, soil physicochemical properties, and beta diversity are visualised by db-RDA. *p* values are denoted by asterisks (* *p* < 0.05, ** *p* < 0.01, *** *p* < 0.001). Results of marginal permutation tests of db-RDA are shown in **Table S1**. **(c)** Relative abundance of the taxa at the phylum level based on 16S rRNA gene amplicon sequencing and metagenome analysis. **(d)** Differences in relative abundance of key genera between the elevated H_2_-treated soils and control soils (0.5 ppmv H_2_ treatment) based on 16S rRNA gene amplicon sequencing and metagenome analysis. The percent values in parentheses refer to the relative abundance of the phylotype in the control soil. Only taxa are shown which significantly increased or decreased in relative abundance by at least 1% in the treatment versus control microcosms, where * indicates *p* < 0.05 (one-way ANOVA with Duncan’s test).

**Table 1.**
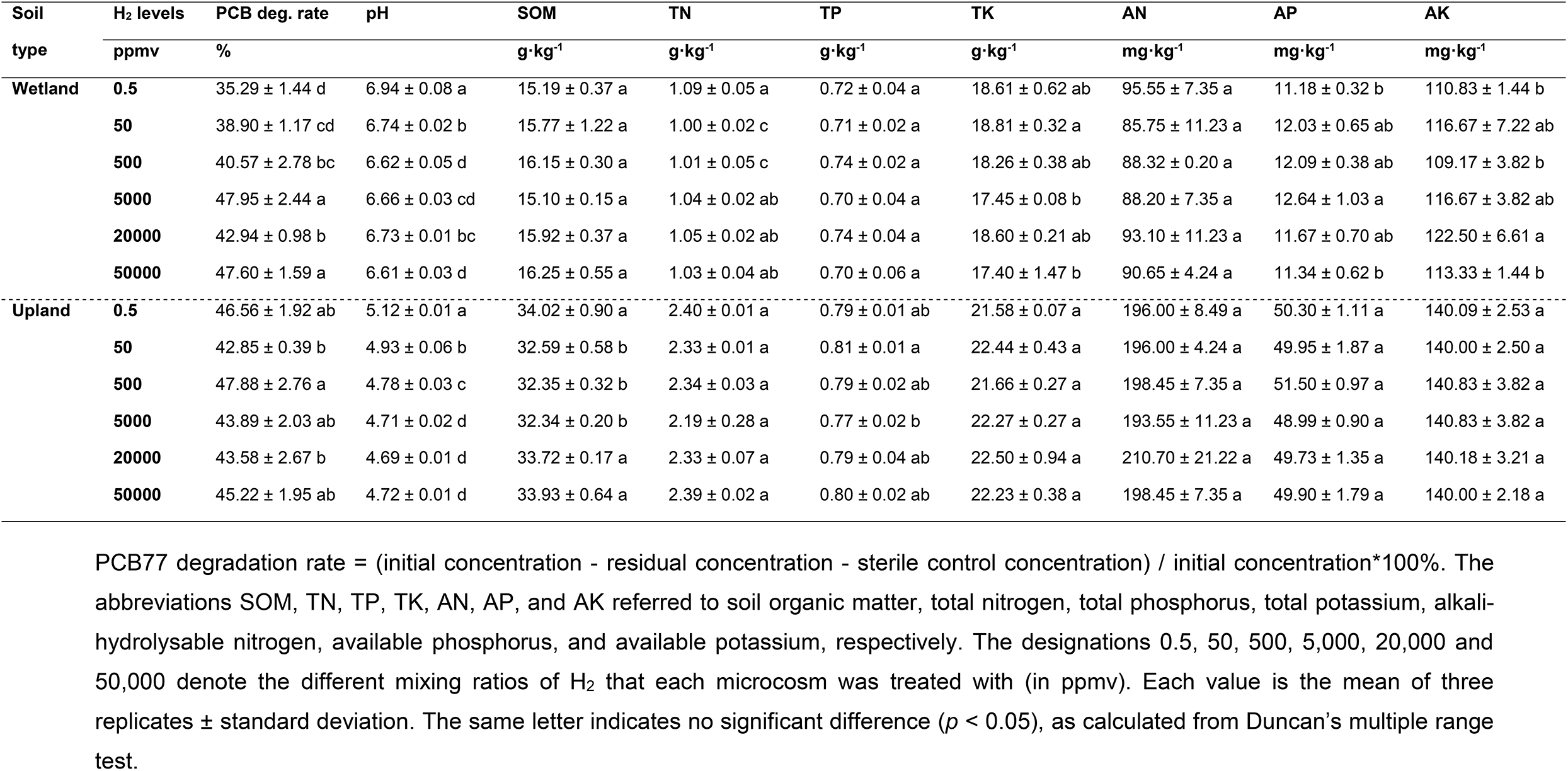
Edaphic properties and PCB degradation rate in different treatments after 84 days.

Microbial community composition was determined using a combination of 16S rRNA gene amplicon sequencing and reconstruction of 16S rRNA gene sequences from metagenomic raw reads via GraftM (metagenomes sequenced for microcosms exposed to 0.5, 20,000, and 50,000 ppmv H_2_ only). Observed phylum-level community composition was comparable between profiles from metagenomic and amplicon sequencing (**Fig. 1c**). In both soils, most community members (>80%) affiliated with six of the globally dominant soil phyla [76, 77], namely Proteobacteria, Firmicutes, Acidobacteriota, Actinobacteriota, Chloroflexota, and Gemmatimonadota (**Fig. 1c**). Significant changes in microbial community composition was observed at both phylum and genus levels in response to H_2_ treatment. In both soils, there decrease in the relative abundance of Firmicutes in the treatment vs control microcosms after 84 days, primarily due to the decline of several Bacilli genera. However, the enriched bacteria strikingly differed between the soils. For the wetland soils, in line with previous observations by Zhang *et al*. [71], there was an enrichment in the phylum Proteobacteria (**Fig.1c & Table S2**). This was driven by significant increases in the relative abundance (by over 1%, *p* < 0.05) of genera such as *Dongia*, and *Noviherbaspirillum* (**Fig. 1d & Table S3**). In contrast, in the upland soils, the most enriched taxa were *Mycobacterium* (Actinobacteriota) and *Candidatus* Koribacter (Acidobacteriota); whereas *Mycobacterium* was a member of the rare biosphere in the control microcosms (0.04% relative abundance), it grew in a dose-dependent manner to become the most abundant genus in the 50,000 ppmv treatments based on both amplicon (5.58%) and metagenomic sequencing (4.86 %) (**Fig. 1d & Table S3**). This observation of a large single-member community shift is remarkably similar to Osborne *et al.*’s RFLP-based inference of the enrichment of actinobacterial taxa, including *Mycobacterium*, following H_2_ exposure in Australian soils [57].

To gain further insight into community responses to elevated H_2_, we assembled and binned the metagenomes, yielding 196 metagenome-assembled genomes (MAGs; **Table S4**). Reconstructed MAGs comprise taxonomically diverse members from a total of two archaeal and 22 bacterial phyla (**Fig. 2** and **Table S4**). In support of 16S rRNA gene amplicon analysis (**Fig. 1d**), most MAGs affiliated with the phyla Proteobacteria (52), Actinobacteriota (30), Acidobacteriota (22), Gemmatimonadota (20), and Chloroflexota (15). We also retrieved a surprising number of Patescibacteria MAGs (17), supporting recent reports that these symbionts can be abundant in oxygenated soils [78, 79]. Based on metagenomic read mapping, there was a significant enrichment of 21 MAGs (12 phyla) in the wetland soil microcosms and 10 MAGs (4 phyla) from in the upland microcosms at high H_2_ treatments, suggesting a complex response at the individual taxon level (**Fig. 2**). In line with the amplicon- and metagenome-based 16S rRNA gene analysis (**Fig. 1 & Table S3**, some MAGs became highly abundant after higher H_2_ treatments, potentially through hydrogenotrophic growth. The most enriched MAGs overall were *Noviherbaspirillum*-affiliated Bin377 in the wetland soil (0.017%, 0.38%, and 0.22% at 0.5, 20,000, and 50,000 ppmv respectively) and *Mycobacterium*-affiliated BinFLU20000R1_4 in the upland soil (0%, 0.59%, and 1.36% at 0.5, 20,000, and 50,000 ppmv respectively) (**Fig. 2**).

**Figure 2.**
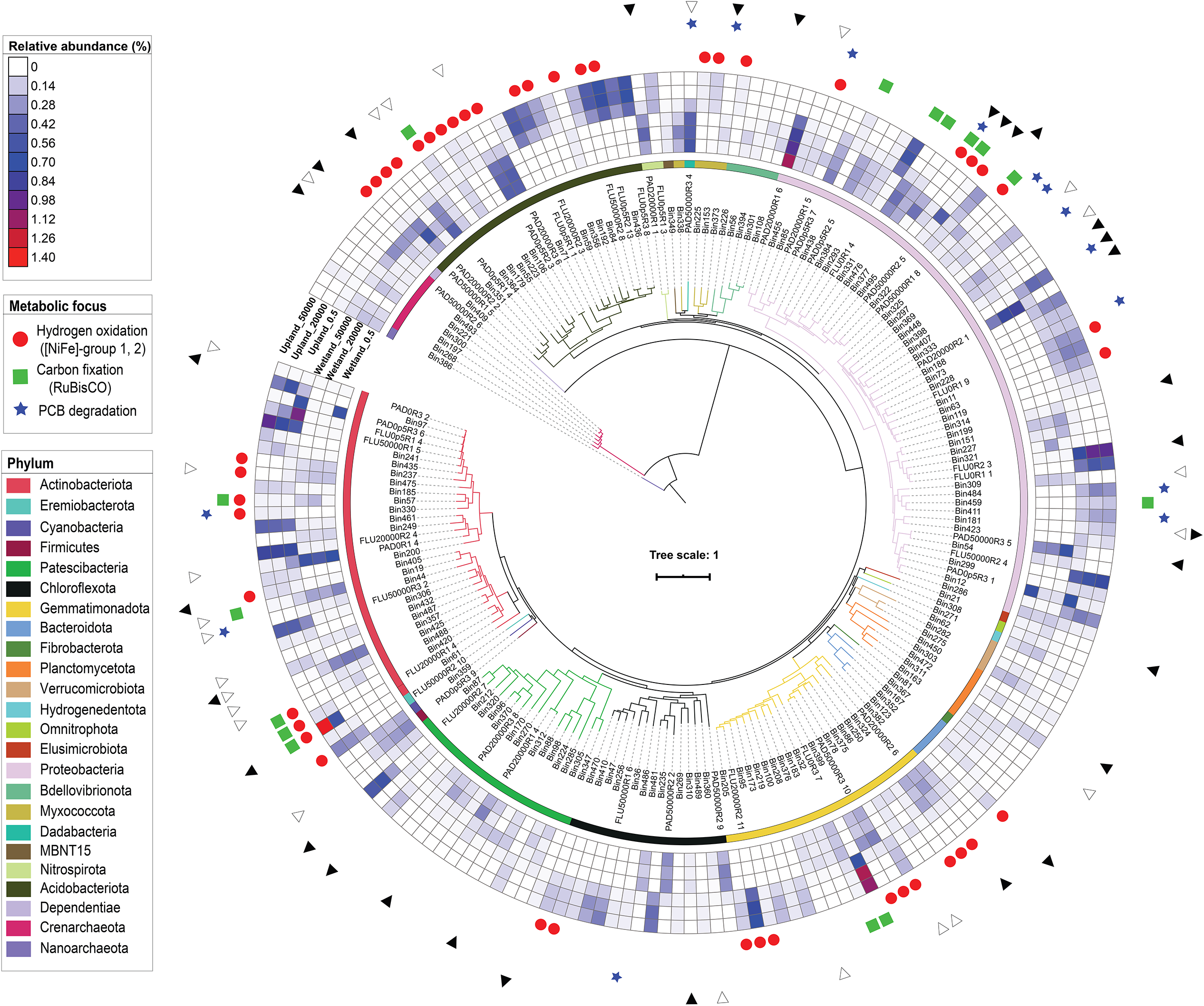
Phylogenetic tree of 196 assembled contaminated soil microbial MAGs. The average abundance of each MAG in the corresponding hydrogen treatments in contaminated soil is shown in the outer circle heatmap. Taxonomy classification at the phylum level is shown in the inner circle across the 196 MAGs spanning 24 phyla. The square indicates MAGs that encode a group 1 or 2 [NiFe]-hydrogenase, the circles indicate MAGs that encode a RuBisCO, and the stars indicate MAGs that encode PCB degradation pathways. The triangle (left triangle, wetland; right triangle, upland) denotes on the diagram those taxa that are significantly changed following H_2_ treatment; Filled symbol indicates significantly enriched MAGs, and symbol with border only indicates significantly decreased MAGs.

### Enriched taxa encode different hydrogenase and RuBisCO lineages known to support hydrogenotrophic growth

We used curated metabolic marker gene databases to annotate the metagenomic short reads and derived MAGs with a focus on H_2_ metabolism and carbon fixation pathways. In agreement with our recent findings in Australian soils [3], most community members were predicted to metabolically versatile with respect to electron donor, electron acceptor, and carbon source preferences (**Fig. 3, Fig. S3, Table S5 & Table S6**). As expected from the community profile (**Fig. 1 & Table S3**), almost all community members encoded markers for aerobic respiration (notably CoxA, CcoN, CydA), with many having the capacity for denitrification (notably NarG, NirK, NorB, NosZ) and hydrogenogenic fermentation (group 3b [NiFe]-hydrogenases) **(Fig. S3)**. With respect to electron donor utilisation, the marker genes for the oxidation of organic compounds (NuoF, SdhA), H_2_ (uptake hydrogenases), carbon monoxide (CoxL), formate (FdhA), and sulfide (Sqr) were abundant in the short reads and widespread in the MAGs. Moreover, there was a widespread capacity for carbon fixation primarily through the Calvin–Benson–Bassham cycle (RbcL) (**Figs. 2 & 3**). As expected, we observed significant increases in the relative abundance of uptake hydrogenases and RuBisCO for both soils in the high H_2_ microcosms, suggesting hydrogenotrophic growth. There were also small but significant changes in the abundance of certain genes involved in aerobic respiration, denitrification, nitrogen fixation, and sulfide, nitrite, and arsenite oxidation. Moreover, in the wetland soils, there was a large enrichment of a gene (BphA) for biphenyl degradation following H_2_ treatment (**Fig. 3 & Table S6**).

**Figure 3.**
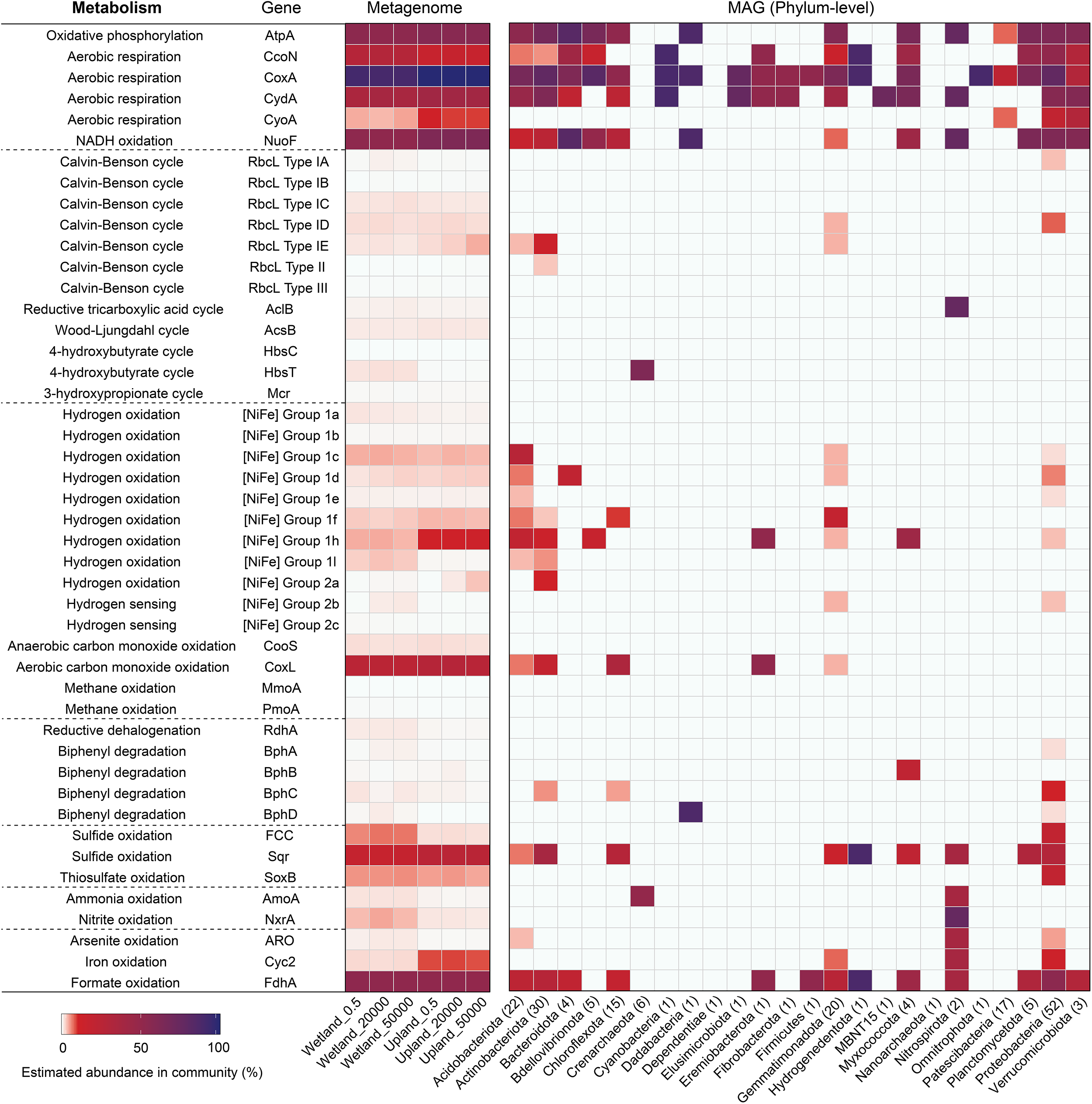
Changes of metabolic potential of the microbial communities in contaminated soils. To infer gene abundance in metagenomes, read counts were normalized to gene length and the abundance of 14 single-copy marker genes; while the abundance of the right heatmap were normalized by predicted MAG completeness.

To gain a deeper insight into the determinants of hydrogenotrophic growth, we built phylogenetic trees to classify the [NiFe]-hydrogenase (**Fig. S4)** and RuBisCO (**Fig. S5**) sequences retrieved from the MAGs based on functionally predictive schemes [80, 81]. Respiratory uptake hydrogenases were encoded by 46 MAGs from nine soil phyla. Many MAGs encoded group 1h [NiFe]-hydrogenases, suggesting they persist in soils by scavenging H_2_ at atmospheric levels; these hydrogenases were present in taxa as diverse as Actinobacteriota (8), Acidobacteriota (6), Proteobacteria (2), Bdellovibrionota (1), Eremiobacterota (1), Gemmatimonadota (1), and Myxococcota (1), in line with recent inferences of a diverse and abundant H_2_ sink in soils [3, 17, 78]. However, a range of taxa also encoded hydrogenases implicated in hydrogenotrophic growth, such as the group 1d and 2a [NiFe]-hydrogenases [37, 82], as well as the functionally enigmatic group 1c and 1f [NiFe]-hydrogenases [3, 45] (**Fig. S4**). Of these, four Actinobacteriota, three Proteobacteria, and one Acidobacteriota MAGs co-encoded uptake hydrogenases with RuBisCO (**Fig. 2**). Seven of these eight MAGs increased in abundance in the H_2_-supplemented soils, including the previously highlighted Bin377 (*Noviherbaspirillum*) and BinFLU20000R1_4 (*Mycobacterium*). This suggests that these bacteria grow hydrogenotrophically by using electrons derived from H_2_ for aerobic respiration and carbon fixation. These metagenomic inferences are supported by previous culture-based studies observing hydrogenotrophic growth in various *Mycobacterium* species [42, 83, 84] and a rice paddy *Noviherbaspirillum* isolate [85]. Thus, the most strongly enriched MAGs in high H_2_ microcosms were among those capable of hydrogenotrophic growth.

The enzyme lineages supporting hydrogenotrophic growth differed between the soils. In the upland soils, the most enriched lineages were Mycobacteriaceae and Pseudonocardiaceae harbouring group 2a [NiFe]-hydrogenases with type IE RuBisCO, such as the *Mycobacterium* MAG (**Fig S4 & S5**). In these soils, the abundance of short reads encoding the group 2a [NiFe]-hydrogenase increased by 17-fold (*p* = 0.0095) and 49-fold (*p* = 0.0028) at H_2_ doses of 20,000 ppmv and 50,000 ppmv respectively. By contrast, in the wetland soils, the most enriched lineages were Burkholderiaceae that encoded group 1d and 2b [NiFe]-hydrogenases together with type IA or IC RuBisCO, including two *Noviherbaspirillum* MAGs. Consistently, in the metagenomic short reads for the wetland soil, there was an increase in relative abundance of the uptake 1d [NiFe]-hydrogenase (1.8-fold, *p* = 0.0034) and the sensory 2b [NiFe]-hydrogenase (15-fold; *p* =0.0025). Based on the precedent of the closely related species *Ralstonia eutropha* (Burkholderiaceae), stimulation of the sensory hydrogenase by elevated H_2_ activates a signal transduction cascade that increases transcription of the uptake hydrogenase and in turn enables hydrogenotrophic growth [86, 87]. Thus, bacteria with the capacity to both sense and oxidise H_2_ can rapidly respond to this energy source becoming available. It should be noted that, while the group 1h and 3b [NiFe]-hydrogenases were the most widespread hydrogenases in both soils overall, their abundance minimally changed in response to H_2_ exposure; this reflects their respective physiological roles in supporting persistence through atmospheric H_2_ oxidation during carbon starvation and fermentative H_2_ production during hypoxia [21, 22, 40]. Consistent with the community composition (**Fig. 1c**), Uptake hydrogenases and other marker genes associated with anaerobic H_2_ oxidation processes (e.g. methanogenesis, acetogenesis, sulfate reduction) were in low abundance in all microcosms.

### Hydrogenotrophic growth of a specific taxon underlies enhanced PCB bioremediation in H_2_-stimulated wetland soils

Of the 86 genes profiled, other than hydrogenases and RuBisCO, the determinants of PCB bioremediation showed the greatest fold change in response to H_2_ supplementation. Based on short reads, in the H_2_-enriched wetland microcosms, we observed a 3.9-fold increase in relative abundance of the genes encoding biphenyl dioxygenase (*bphA*). No equivalent enrichment was observed in the upland soil, by contrast. At the MAG level, 14 MAGs encoded enzymes for biphenyl oxidation to benzoate (Proteobacteria, Myxocococcota, Chloroflexota, Dadabacteria) (**Fig. S6 & Table S7**). Of these, the metabolic capabilities of four MAGs (>90% completeness, <5% contamination) are depicted in **Fig. 4a**. Five of these MAGs increased in abundance at elevated H_2_ concentrations in wetland soils, including the aforementioned Bin377 (*Noviherbaspirillum*), which was the sole MAG encoding the *bphA* gene.

**Figure 4.**
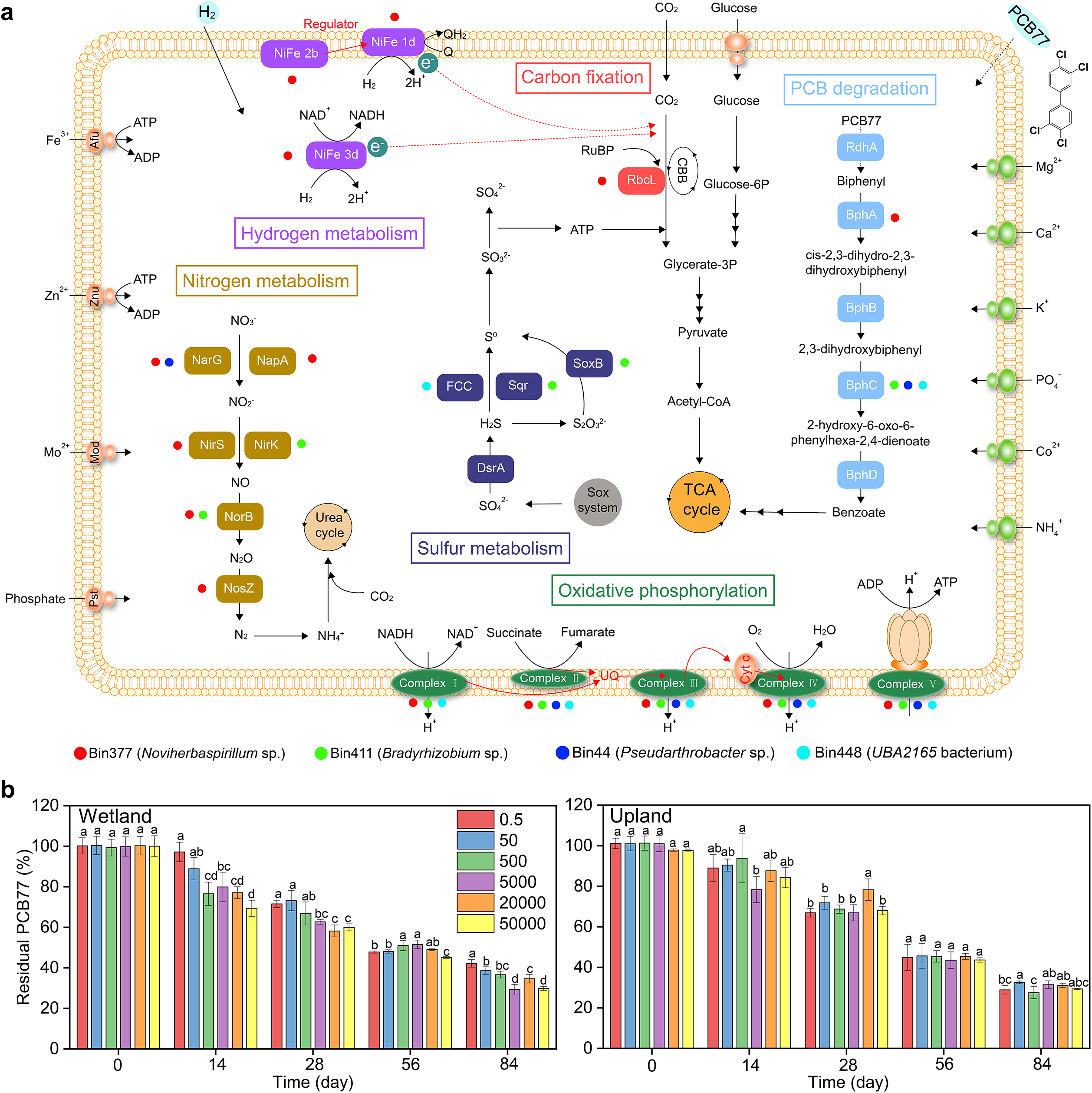
Effect of hydrogen on PCB degradation genes and activities in contaminated soils. **(a)** Metabolic pathways in four MAGs (with > 90% completeness and < 5% contamination) predicted to mediate PCB degradation. Predicted proteins in the figure are listed in **Table S7 and Table S11**. **(b)** Changes of residual PCB77 concentrations in the wetland and upland soils during the incubation period.

Consistent with these observations, we observed divergent effects of elevated H_2_ on PCB77 biodegradation in the two soils (**Fig. 4b**). After 84 days, the degradation rate of PCB77 in the wetland soil at elevated H_2_ concentrations (5000-50000 ppmv) were significantly promoted by 7.65 to 12.66% compared with the 0.5 ppmv (*p* < 0.05, **Table 1 & Fig. 4b**). By contrast, there were no significant promotion of PCB77 degradation in the upland soil at elevated H_2_ concentrations during the experimental period (**Table 1 & Fig. 4b**). Altogether, these findings provide a mechanistic rationale for our previous observations that PCB bioremediation is enhanced both by nitrogen fixation (resulting in endogenous H_2_ production) and endogenous H_2_ addition [60, 61]; the enrichment of hydrogenotrophic Burkhoderiaceae and likely other taxa encoding biphenyl oxidation genes enhances bioremediation primarily through indirect effects.

## Conclusions

This study demonstrates that phylogenetically and physiologically diverse H_2_-oxidising bacteria reside in soils. Whereas most of these bacteria are organoheterotrophs predicted to persist on trace concentrations of H_2_, a few community members are facultative autotrophs that grow on this gas when available in elevated concentrations. In our microcosms, the bacterial, hydrogenase, and RuBisCO lineages that were enriched in response to H_2_ availability strikingly differed between the soils, in a way that reflected their native community composition. In the upland soil, the *Mycobacterium* MAG and several other lineages emerged from the rare biosphere to become dominant community members in the upland soils, whereas in the wetland soil a *Noviherbaspirillum* MAG possessing two hydrogenases likely sensed and rapidly consumed high concentrations of H_2_. These findings in turn provide a holistic community context to previous culture-based investigations on hydrogenotrophic Actinobacteriota and Proteobacteria. Moreover, these observations findings reconcile the seemingly divergent findings of earlier RFLP-based studies in this area [57, 71], though are less compatible with certain recent reports [67, 72, 74]. Overall, we found that H_2_ supplementation did not profoundly affect microbial community abundance, diversity, or capabilities. However, it can be expected that enrichment of hydrogenotrophic taxa will have various effects on biogeochemical activities, as reflected by the increased genetic capacity and biochemical activity for PCB biodegradation captures in the wetland. Extending these findings, we predict that hydrogen emissions from natural or anthropogenic sources would select for the growth of facultative hydrogenotrophs, though the lineages stimulated are likely to greatly vary between soils.

## Methods

### Experimental soils

The top 20 cm of the soil profile of two agricultural soils were sampled for the microcosm experiments. An anthrosols soil, known to have a high capacity for pollutant remediation [88], was sampled from long-term paddy wetland field experimental station of the Chinese Academy of Sciences located at Changshu, Jiangsu province (31°33′N, 120°38′E). A flavo-aquic soil was sampled at a long-term abandoned meadow upland field, formerly highly polluted by PCBs, from Taizhou, Zhejiang province (28°31′N, 121°22′E). Both soils were air-dried in the laboratory and then passed through a 10-mesh screen to remove roots and large particles before the preparation of soil microcosms. PCB levels in both soils are within permittable levels, with total concentrations of 21 PCBs below 60 μg kg^−1^ and no PCB77 detected in either soil. Detailed soil properties are listed in **Table S8**.

### Microcosm setup and sampling

Prior to the microcosm setup, each soil was adjusted to a moisture of 10% (w/w) and preconditioned at 30 °C for one week. In order to monitor effects of H_2_ supplementation on soil microbial communities and bioremediation, PCB77 was also added to both soils to a final concentration of 1 mg kg^−1^. Specifically, 15 mL of PCB77 stock solution (100 mg L^−1^ in acetone) was added to 150 g soil (dry weight) to achieve a concentration of 10 mg kg^−1^; the soils were placed in a fume hood overnight to evaporate the acetone and they were then thoroughly mixed with 1350 g uncontaminated soil. Thereafter, approximately 10 g of soil (dry weight) was placed in a 120 mL serum bottle and adjusted to a water content of 30 % (w/w) with sterilized water. The serum bottles were sealed with butyl rubber stoppers. The bottles were flushed with synthetic air (360 ppmv CO_2_ and 21 % O_2_ balanced with N_2_; 55th Research Institute, China Electronics Technology Group Corporation) for 30 s and then an appropriate volume of synthetic air was withdrawn. A defined volume of ultra-pure H_2_ (99.9999%; 55th Research Institute, China Electronics Technology Group Corporation) gas was injected to obtain six initial headspace mixing doses of H_2_ (0.5, 50, 500, 5,000, 20,000, and 50,000 ppmv). The 0.5 ppmv vials served as controls, given they reflect ambient H_2_ concentrations, whereas the five other vials served as treatment groups with elevated H_2_ levels. A sterile control (autoclaving at 121 °C, 1 h three times) was also setup to exclude the factors of soil adsorption of PCB77 (**Fig. S7**). Each day, the control, treatment, and sterile control serum bottles were flushed with synthetic air and then the initial concentrations of H_2_ were re-established as described above, thereby providing a regular H_2_ supply. All treatments were set up in triplicate and incubated at 30 °C in the dark for 84 days. Samples were taken on days 0, 14, 28, 56 and 84 for DNA extraction and PCB77 quantification.

### Analysis of physicochemical properties and PCB77 contents

To determine physicochemical properties and quantify PCB77 levels, freeze-dried soil samples were sieved through a 60-mesh screen to obtain a homogeneous matrix.

The pH, soil organic matter content (SOM), total nitrogen (TN), total phosphorus (TP), total potassium (TK), alkali-hydrolyzable nitrogen (AN), available phosphorus (AP), and available potassium (AK) were measured as previously described [89]. Briefly, soil pH was determined in a soil/water suspension (1:2.5) using pH meter; SOM was measured by the K_2_Cr_2_O_7_-H_2_SO_4_ oxidation method; TN was determined by Kjeldahl digestion; TP and TK were determined by molybdenum-blue colorimetry and flame photometry respectively after HF-HClO_4_ treatment; AN was assayed by alkali-hydrolyzed diffusion method; AP was determined by sodium bicarbonate extraction and molybdenum blue colorimetry; and AK was detected by ammonium acetate extraction and subsequent flame photometer analysis.

Soil PCB77 was extracted as described by Huang et al [90]. PCB 77 concentrations were detected by GC7890 gas chromatograph (Agilent Technologies, Santa Clara, CA) equipped with a HP5 column (30 m × 0.32 mm × 0.25 μm). The recovery rates for all the samples and detection limit of the GC method for PCB77 ranged from 87 to 102 % and 2.53 to 5.75 μg kg^−1^ respectively.

### DNA extraction

Community DNA was extracted from fresh soil using the FastDNA spin kit for soil (MP Biomedicals, Santa Ana, CA) following the manufacturer’s instructions. Sample DNA integrity was examined by electrophoresis on a 0.8 % agarose gel. Sample DNA quantity and purity were determined with a Nanodrop ND-2000 UV-Vis spectrophotometer (NanoDrop Technologies, Wilmington, DE) and using Quant-iT PicoGreen fluorescence (Thermo Fisher, Waltham, MA). The DNA samples were stored at −80 °C before use.

### Quantitative PCR assays

Quantitative PCR (qPCR) was used to quantify the copy number of 16S rRNA genes for (i) whole bacterial and archaeal community and (ii) anaerobic PCB-degrading genus *Dehalobacter* [91]. PCR amplification was performed using the primer sets 515F/907R and Dhb-477F/ 647R **(Table S9)** with cycling conditions as previously described [92]. The qPCR reactions were conducted for each soil DNA extract in triplicate. Standard curves were established using a linear PCR product by a 10-fold serial dilution of plasmid DNA that contained the target fragment. The amplification efficiencies were 97.5-99.8%.

### 16S rRNA gene amplicon sequencing and analysis

The V4-V5 region of the 16S rRNA gene was used to determine the composition of the soil microbial communities with universal prokaryotic primer sets 515F/907R **(Table S9)** for 36 samples at 84 days (0.5, 50, 500, 5000, 20000, and 50000 ppmv H_2_ treatment). Amplification of the 16S rRNA gene target was performed according to the manufacturer’s instructions (Illumina). The amplicons were sequenced by the Majorbio Company (Shanghai, China) following the manufacturer’s instructions with a MiSeq PE300 platform (Illumina, San Diego, CA, USA).

The resulting raw reads were processed on the QIIME2 platform (version 2020.02) using the DADA2 pipeline to resolve exact amplicon sequence variants (ASVs) [93, 94].The taxonomy of each 16S rRNA gene sequence was analysed using RDP classifier algorithm (http://rdp.cme.msu.edu/) against Silva (SSU132) 16S rRNA gene database and the Unite (Release 6.0) database using a confidence threshold of 70% [95]. Alpha diversity (including observed richness, Chao1 richness, and Shannon index) and beta diversity (Bray Curtis) were calculated with mothur (version v.1.30.1, collect single command) and QIIME2 with the default parameters, respectively [96]. The relationships between beta diversity and environmental variables were displayed through distance-based redundancy analyses (db-RDA) based on Bray-Curtis distance (R: vegan package, version 4.0.3). A one-way analysis of variance (ANOVA) to test for significant differences in community structure between different H_2_ concentration treatments.

### Metagenomic sequencing, assembly and binning

Based on the results of 16S rRNA gene amplicon sequencing, the samples from higher H_2_ concentrations treatments (20,000 and 50,000 ppmv) showing obvious succession of bacterial communities with 0.5 ppmv at 84 days were subject to metagenomic analysis. The extracted DNA was sheared into approximately 400 bp fragments using a Covaris M220 shaker (Gene Company Limited, China). The metagenomic libraries were prepared using the NEXTFLEX Rapid DNA-Seq Kit (PerkinElmer Bioo Scientific, USA). Paired-end sequencing was performed on the Illumina HiSeq 4000 platform (Illumina Inc., San Diego, CA, USA) at Shanghai Majorbio Bio-pharm Technology Co., Ltd. About six billion base pairs (~6 Gbp) of DNA sequences were generated for each sample **(Table S10)**. To explore microbial composition of each sample, taxonomic assignments of raw reads were assigned using GraftM [97] together with Silva (SSU132) 16S rRNA gene database [95].

Raw reads were quality-controlled using Read_QC module in the metaWRAP pipeline [98]. The quality-controlled metagenomes were individually assembled and co-assembled using MEGAHIT v1.1.3 (default parameters) [99]. The resulting assemblies were binned using the binning module within the metaWRAP pipeline (--metabat2 --maxbin2 --concoct for individual assembly; --metabat2 for co-assembly). For each assembly, the three bin sets were then consolidated into a final bin set with the bin_refinement module of metaWRAP (-c 50 -x 10 options). The final bin sets from both individual assemblies and co-assembly were aggregated and de-replicated using dRep v2.5.4 [100] at 95% average nucleotide identity (-comp 50 -con 10 options). The quality (completeness and contamination) of MAGs was assessed with CheckM [101].

The taxonomy of each MAG was temporally assigned using GTDB-Tk [102] (GTDB R04-RS89 database). The relative abundance of each MAGs was calculated with CoverM (https://github.com/wwood/CoverM) as previously described [103]. The taxonomic classification, size, completeness, contamination, strain heterogeneity, and N50 of recovered MAGs are summarized in **Table S4**.

### Functional annotation of reads and MAGs

For functional annotation of quality-filtered reads with lengths over 140 bp, metabolic marker genes covering the major pathways associated with hydrogen cycling, carbon fixation, oxidative phosphorylation, anaerobic PCB degradation (reductive dehalogenation), and the cycling of nitrogen compounds, sulfur compounds, methane, and carbon monoxide were searched as previously described [78]. DIAMOND *blastx* mapping [104] was performed with a query coverage threshold of 80% for all databases, and a percentage identity threshold of 50%, except for group 4 [NiFe]-hydrogenases, [FeFe]-hydrogenases, CoxL, AmoA, and NxrA (all 60%), PsaA (80%), PsbA and IsoA (70%), and HbsT (75%). These reads were then transformed to per kilobase per million (RPKM) [105]. Functional annotation of putative amino acid sequences involved in aerobic PCB degradation (biphenyl degradation, including BphA, BphB, BphC and BphD) was searched as described in the **supplementary note**. The gene abundance in the microbial community was then estimated by the method according to Ortiz et al., [78]. Briefly, 14 universal single copy ribosomal marker genes were also transformed to RPKM and gene abundance in the microbial community was calculated by dividing the read count for the gene (in RPKM) by the mean of the read counts of the 14 universal single copy ribosomal marker genes (in RPKM). For each MAG, genes were called by Prodigal (-p meta) [106]. Genes involved metabolic functions as described above were carried out using DIAMOND *blastp* with a minimum percentage identity of 60% (NuoF), 70% (AtpA, ARO, YgfK) or 50% (all other databases) [78], while genes involved in biphenyl degradation were annotated against KEGG database using GhostKOALA [107].

### Phylogenetic analysis

For phylogenetic tree construction of MAGs, ribosomal protein sequences generated from CheckM were extracted and aligned using MAFFT [108]. Gaps in the alignment were removed and the ribosomal protein alignment concatenated as described previously [109]. RAxML webserver (https://www.phylo.org/) was used to construct the phylogenetic tree with the parameters: raxmlHPC-HYBRID -f a -n result -s input -c 25 -N 160 -p 12345 -m PROTCATLG -x 12345, with the output file uploaded to iTOL for visualization [110].

For amino acid sequences of the group 1, 2 [NiFe]-hydrogenase and ribulose 1,5-bisphosphate carboxylase/oxygenase (RuBisCO) large subunit (RbcL), sequences were aligned using the ClustalW algorithm included in MEGA7 [111]. Their maximum-likelihood phylogenetic trees were constructed using the JTT matrix-based model, and was bootstrapped with 50 replicates and midpoint-rooted.

## Supplementary information

Supplementary information (Figure S1-S7; Table S1-S11; Supplementary text and Supplementary Tables (xlsx)) accompanies this paper.

## Acknowledgments

This study is funded by the National Key Research and Development Program of China (2019YFC1803700) and the National Natural Science Foundation of China (Grant Nos. 41671327). C.G. is supported by an ARC DECRA Fellowship (DE170100310) and an NHMRC EL2 Fellowship (APP1178715). X.D. is supported by National Natural Science Foundation of China (Grant no. 41906076) and the Fundamental Research Funds for the Central Universities (Grant no. 19lgpy90).

## Author contributions

Y.X., Y.T., and Y.L. conceived and supervised this study. Y.X., Y.T., and X.W. designed and performed experiments. Y.X., X.D., Y.T., C.Z., and C.G. were responsible for meta-omic analysis. Y.X., X.D., and C.G. analysed data. L.Z. and W.R. provided critical comments on this study. Y.X., Y.T., X.D., and C.G. wrote the paper with input from all authors.

## Data availability statement

The 16S rRNA gene amplicon sequences have been deposited in the Sequence Read Archive (SRA) of the NCBI with accession number PRJNA639898. The metagenomes have been deposited in the NCBI SRA with accession number PRJNA640224.

## Abbreviation list

SOM: soil organic matter content
TN: total nitrogen
TP: total phosphorus
TK: total potassium
AN: alkali-hydrolyzable nitrogen
AP: available phosphorus
AK: available potassium.

## Ethics approval and consent to participate

Not applicable.

## Consent for publication

Not applicable.

## Statement

We confirm we have included a statement regarding data and material availability in the declaration section of our manuscript.

