## Supplementary for "Distinct hydrogenotrophic bacteria are stimulated by elevated H_2_ levels in upland and wetland soils"

### Supplementary information

#### Supplementary note: Functional annotation of aerobic PCB degradation

To improve the reliability and quality of subsequent analysis, the raw sequence data was processed with the following steps: (i) SeqPrep (https://github.com/jstjohn/SeqPrep, default parameters) was used to strip the adapter sequences from the 3′ and 5′ ends of the paired-end Illumina reads; (ii) Sickle (https://github.com/najoshi/sickle) was used to trim low-quality reads (quality value < 20 or contain N bases) and short reads (<50 bp). Following quality filtering, an average of 98.71% high quality reads was obtained from all samples. Multiple_Megahit was used to combine contigs with minimum lengths longer than 300 bp [1]. In brief, the high quality reads of each sample were assembled *de novo* using Megahit and then unmapped reads were also assigned under default parameters through Megahit [2]. Finally, these contigs were combined and used as final assembly results for gene prediction. For assembly, one additional metagenome (without $H_2$ treatments) sequenced using the same method was also included. Open reading frames (ORFs) were predicted using MetaGene [3]. ORFs (>100 bp) were translated to amino acid sequences using the NCBI translation table. All predicted genes (under default parameters: 95% identity and 90% coverage) to references were clustered using CD-HIT, and the longest sequences from each cluster was chosen as representative sequence to construct a non-redundant gene catalogue [4]. Quality-filtered reads were mapped to these representative sequences with 95% identity using SOAPaligner to determine the gene abundances in each sample [5]. Functional annotation of putative amino acid sequences involved in aerobic PCB degradation (including BphA, BphB, BphC and BphD), which were translated from the gene catalogue, was carried out through DIAMOND blastp search (Version 2.2.28+) against the Kyoto Encyclopedia of Genes and Genomes (KEGG) database (cutoffs: e-value 1e-5, identity: 60%) [6]. The abundance of each gene was converted to per kilobase per million (RPKM) described in previous study [7] as following:

$$RPKM_i = \frac{R_i * 10^6}{L_i * \sum_1^n R_i}$$

$PRKM_i$: The relative abundance of gene i in sample S.

$R_i$: The number of mapped reads which gene i could be detected in sample S.

$L_i$: The length of gene i.

**Table S1. Results of marginal permutation tests of db-RDA for Figure 1b.**

| Predictor | Wetland | | | | Upland | | | |
|---|---|---|---|---|---|---|---|---|
| | CAP1 | CAP2 | $R^2$ | *P* value | CAP1 | CAP2 | $R^2$ | *P* value |
| $H_2$ | 0.1260 | 0.9920 | 0.819 | **0.001** | -0.4905 | -0.8714 | 0.950 | **0.001** |
| pH | -0.8103 | -0.5860 | 0.629 | **0.001** | 0.2820 | 0.9594 | 0.320 | **0.047** |
| SOM | -0.3874 | 0.9219 | 0.151 | 0.317 | -0.8232 | -0.5678 | 0.266 | 0.100 |
| TN | -0.5936 | -0.8048 | 0.137 | 0.314 | -0.2112 | -0.9774 | 0.040 | 0.505 |
| TP | -0.8372 | 0.5469 | 0.033 | 0.773 | 0.8925 | -0.4511 | 0.068 | 0.636 |
| TK | -0.9147 | -0.4041 | 0.375 | **0.027** | -0.5263 | -0.8503 | 0.070 | 0.589 |
| AN | -0.9992 | 0.0390 | 0.298 | 0.068 | 0.2056 | -0.9786 | 0.102 | 0.320 |
| AP | 0.9158 | -0.4017 | 0.411 | **0.020** | 0.9966 | -0.0818 | 0.287 | 0.062 |
| AK | 0.5036 | 0.8639 | 0.103 | 0.426 | 0.9347 | -0.3555 | 0.238 | 0.104 |

The abbreviations SOM, TN, TP, TK, AN, AP, and AK refer to soil organic matter, total nitrogen, total phosphorus, total potassium, alkali-hydrolyzable nitrogen, available phosphorus, and available potassium, respectively.

**Table S2. Results of one-way analysis of variance (ANOVA) between treatments for Figure 1c.**

| Soil type | Taxonomy | 16S rRNA gene amplicon | | | | | | | | | | | | Metagenome | | | | | |
|---|---|---|---|---|---|---|---|---|---|---|---|---|---|---|---|---|---|---|---|
| | | 0.5 | | 50 | | 500 | | 5,000 | | 20,000 | | 50,000 | | 0.5 | | 20,000 | | 50,000 | |
| | | SD | Sig. | SD | Sig. | SD | Sig. | SD | Sig. | SD | Sig. | SD | Sig. | SD | Sig. | SD | Sig. | SD | Sig. |
| Wetland | Acidobacteria | 0.35 | ab | 0.37 | ab | 0.70 | ab | 0.55 | ab | 0.45 | b | 1.84 | a | 0.49 | b | 0.30 | a | 2.73 | a |
| | Actinobacteria | 0.35 | a | 0.67 | a | 0.17 | a | 0.87 | a | 0.66 | a | 0.42 | a | 0.26 | a | 0.42 | a | 0.74 | a |
| | Armatimonadetes | 0.15 | bc | 0.07 | bc | 0.05 | bc | 0.31 | ab | 0.17 | c | 0.18 | a | 0.03 | a | 0.03 | b | 0.38 | b |
| | Bacteroidetes | 0.44 | a | 0.35 | a | 0.39 | a | 0.74 | a | 0.50 | a | 0.18 | a | 0.28 | b | 0.16 | b | 0.11 | a |
| | Chloroflexi | 0.66 | a | 0.73 | a | 1.13 | a | 1.37 | a | 1.49 | a | 0.61 | a | 0.11 | a | 0.31 | a | 0.50 | a |
| | Cyanobacteria | 0.12 | a | 0.17 | a | 0.54 | a | 0.46 | a | 0.20 | a | 0.07 | a | 0.09 | a | 0.34 | a | 0.40 | a |
| | Dadabacteria | 0.40 | a | 0.03 | a | 0.16 | a | 0.28 | a | 0.14 | a | 0.24 | a | 0.00 | a | 0.06 | a | 0.11 | a |
| | Firmicutes | 2.77 | a | 2.21 | ab | 3.31 | b | 5.55 | c | 1.27 | ab | 1.21 | b | 0.94 | a | 0.95 | a | 2.36 | a |
| | Gemmatimonadetes | 0.13 | a | 0.95 | a | 1.09 | a | 1.46 | a | 0.10 | a | 0.34 | a | 0.30 | a | 0.15 | a | 0.81 | a |
| | Patescibacteria | 0.11 | a | 0.04 | a | 0.37 | a | 0.28 | a | 0.18 | a | 0.38 | a | 0.36 | a | 0.20 | a | 0.18 | a |
| | Planctomycetes | 0.45 | b | 0.28 | a | 0.40 | a | 0.29 | a | 0.02 | a | 0.21 | a | 0.25 | a | 0.20 | b | 0.47 | b |
| | Proteobacteria | 2.63 | d | 0.55 | cd | 0.93 | ab | 4.83 | a | 0.76 | c | 1.98 | bc | 1.66 | a | 0.56 | a | 0.43 | a |
| | Rokubacteria | 0.08 | b | 0.06 | b | 0.22 | a | 0.13 | a | 0.09 | b | 0.08 | ab | 0.08 | a | 0.19 | a | 0.15 | a |
| | Others | 0.25 | b | 0.15 | ab | 0.51 | ab | 0.61 | a | 0.31 | ab | 0.52 | ab | 0.32 | a | 0.38 | a | 1.28 | a |
| Upland | Acidobacteria | 1.71 | ab | 2.14 | ab | 5.19 | b | 1.20 | ab | 1.86 | ab | 0.97 | a | 0.33 | a | 0.73 | a | 1.59 | a |
| | Actinobacteria | 3.76 | ab | 0.66 | b | 2.91 | b | 0.71 | b | 1.36 | b | 0.57 | a | 0.71 | ab | 0.72 | b | 2.28 | a |
| | Armatimonadetes | 0.01 | a | 0.22 | a | 0.17 | a | 0.09 | a | 0.24 | a | 0.23 | a | 0.12 | a | 0.22 | a | 0.07 | a |

| | | | | | | | | | | | | | | | | | | |
|---|---|---|---|---|---|---|---|---|---|---|---|---|---|---|---|---|---|---|
| Bacteroidetes | 0.06 | c | 0.11 | bc | 0.06 | c | 0.10 | bc | 0.17 | ab | 0.22 | a | 0.72 | a | 1.40 | a | 0.40 | a |
| Chloroflexi | 0.41 | a | 0.44 | a | 3.36 | a | 0.68 | a | 0.70 | a | 0.31 | a | 0.99 | a | 1.35 | a | 1.42 | a |
| Cyanobacteria | 0.03 | ab | 0.13 | b | 0.14 | b | 0.07 | ab | 0.10 | ab | 0.17 | a | 0.38 | a | 0.60 | a | 0.34 | a |
| Dadabacteria | 0.00 | a | 0.00 | a | 0.00 | a | 0.00 | a | 0.00 | a | 0.01 | a | 0.18 | a | 0.07 | a | 0.20 | a |
| Firmicutes | 0.51 | a | 1.58 | a | 4.73 | abc | 2.35 | ab | 2.30 | bc | 1.71 | c | 0.24 | a | 0.37 | b | 0.24 | b |
| Gemmatimonadetes | 0.25 | bc | 0.32 | a | 0.33 | c | 0.16 | ab | 0.40 | a | 0.18 | a | 1.05 | b | 0.49 | a | 1.31 | ab |
| Patescibacteria | 0.00 | b | 0.04 | ab | 0.00 | b | 0.03 | ab | 0.02 | b | 0.04 | ab | 0.04 | b | 0.07 | a | 0.29 | a |
| Planctomycetes | 0.47 | a | 0.21 | a | 0.66 | a | 0.39 | a | 0.69 | a | 0.34 | a | 0.02 | a | 0.06 | a | 0.30 | a |
| Proteobacteria | 1.46 | a | 1.52 | a | 3.77 | a | 2.27 | a | 1.47 | a | 1.32 | a | 0.01 | a | 0.01 | a | 0.01 | a |
| Rokubacteria | 0.00 | a | 0.00 | a | 0.00 | a | 0.00 | a | 0.00 | a | 0.01 | a | 0.01 | a | 0.04 | a | 0.01 | a |
| Others | 0.49 | a | 0.14 | a | 0.61 | a | 0.15 | a | 0.16 | a | 0.13 | a | 0.19 | a | 0.19 | a | 0.23 | a |

The designations 0.5, 20,000, and 50,000 denote the different mixing ratios of $H_2$ that each microcosm was treated with (in ppmv).

**Table S8. Chemical properties of the two types of soils (Wetland and Upland) used in this study.**

| Parameters (units) | Wetland | Upland |
|---|---|---|
| pH | 7.01 ± 0.01 | 5.44 ± 0.07 |
| SOM ($g·kg^{-1}$) | 19.6 ± 0.40 | 39.70 ± 2.43 |
| TN ($g·kg^{-1}$) | 1.10 ± 0.04 | 2.68 ± 0.10 |
| TP ($g·kg^{-1}$) | 0.71 ± 0.02 | 1.04 ± 0.02 |
| TK ($g·kg^{-1}$) | 19.8 ± 0.13 | 19.63 ± 0.29 |
| AN ($mg·kg^{-1}$) | 69.8 ± 3.68 | 191.10 ± 3.68 |
| AP ($mg·kg^{-1}$) | 8.89 ± 0.47 | 54.11 ± 0.90 |
| AK ($mg·kg^{-1}$) | 113.3 ± 1.44 | 118.33 ± 1.44 |
| CEC ($cmol·kg^{-1}$) | 18.7 ± 0.20 | 18.37 ± 0.81 |
| ∑ 21 PCBs ($μg·kg^{-1}$) | 22.61 ± 8.15 | 40.15 ± 9.56 |
| PCB77 ($μg·kg^{-1}$) | N.D | N.D |

∑ 21 PCBs represent the total concentration of the PCB8, PCB18, PCB28, PCB44, PCB52, PCB66, PCB77, PCB101, PCB105, PCB118, PCB126, PCB128, PCB138, PCB153, PCB170, PCB180, PCB187, PCB195, PCB200, PCB206 and PCB209. The abbreviations SOM, TN, TP, TK, AN, AP, AK, and CEC refer to soil organic matter, total nitrogen, total phosphorus, total potassium, alkali-hydrolyzable nitrogen, available phosphorus, available potassium, and cation exchange capacity, respectively. N.D, not detected.

**Table S9. Primers used for real-time quantitative PCR.**

| Target gene | Primer | Primer sequence 5'-3' | Length | Reference |
|---|---|---|---|---|
| *16SrRNA* | 515F | GTGCCAGCMGCCGCGG | 392 bp | 8 |
| | 907R | CCGTCAATTCMTTTRAGTTT | | |
| *Dehalobacter spp. 16S rRNA* | Dhb 477F | GATTGACGGTACCTAACGAGG | 169 bp | 9 |
| | Dhb 647R | TACAGTTTCCAATGCTTTACGG | | |

**Table S10. Statistics of metagenomic data from quality-filtered unassembled reads and after assembly. Data are shown for the three replicates.**

| Soil type | | 0.5_1 | 0.5_2 | 0.5_3 | 20000_1 | 20000_2 | 20000_3 | 50000_1 | 50000_2 | 50000_3 |
|---|---|---|---|---|---|---|---|---|---|---|
| Wetl and | Number of reads | 46,226,230 | 45,356,086 | 45,686,828 | 46,958,156 | 47,548,956 | 45,712,536 | 46,039,228 | 45,675,672 | 46,202,486 |
| | Number of PE reads | 45,442,324 | 44,664,754 | 44,909,854 | 46,437,048 | 47,052,420 | 45,137,000 | 45,583,598 | 45,198,972 | 45,782,106 |
| | Raw base data (bp) | 6,980,160,73 0 | 6,848,768,98 6 | 6,898,711,02 8 | 7,090,681,55 6 | 7,179,892,35 6 | 6,902,592,93 6 | 6,951,923,42 8 | 6,897,026,47 2 | 6,976,575,38 6 |
| | Number of contigs | 491,369 | 482,937 | 431,111 | 501,141 | 537,512 | 503,043 | 480,800 | 463,252 | 478,483 |
| | Total length (bp) | 313,965,468 | 306,303,549 | 271,231,251 | 318,372,598 | 330,147,128 | 309,458,043 | 299,797,950 | 277,558,303 | 285,784,764 |
| | Max contig length (bp) | 68,098 | 68,098 | 368,185 | 68,098 | 67,991 | 68,098 | 68,098 | 68,098 | 75,490 |
| | Min contig length (bp) | 300 | 300 | 300 | 300 | 300 | 300 | 300 | 300 | 300 |
| | N50 (bp) | 671 | 668 | 666 | 656 | 631 | 633 | 634 | 604 | 601 |
| | N90 (bp) | 349 | 348 | 344 | 348 | 346 | 346 | 345 | 341 | 341 |
| Upl and | Number of reads | 45,541,054 | 45,924,622 | 45,413,340 | 45,952,508 | 46,805,326 | 46,352,782 | 46,832,838 | 46,224,704 | 46,094,660 |
| | Number of PE reads | 44,894,976 | 45,181,452 | 44,735,340 | 45,488,544 | 46,001,684 | 45,850,200 | 46,234,018 | 45,673,962 | 45,578,604 |
| | Raw base data (bp) | 6,876,699,15 4 | 6,934,617,92 2 | 6,857,414,34 0 | 6,938,828,70 8 | 7,067,604,22 6 | 6,999,270,08 2 | 7,071,758,53 8 | 6,979,930,30 4 | 6,960,293,66 0 |
| | Number of contigs | 551,751 | 576,545 | 578,965 | 585,206 | 613,548 | 612,815 | 595,618 | 611,427 | 614,352 |
| | Total length (bp) | 401,662,178 | 417,147,297 | 424,850,940 | 406,547,449 | 438,395,105 | 438,718,815 | 403,395,571 | 406,927,187 | 411,168,616 |
| | Max contig length (bp) | 116,264 | 98,302 | 78,727 | 75,572 | 64,473 | 57,442 | 113,729 | 69,711 | 75,576 |
| | Min contig length (bp) | 300 | 300 | 300 | 300 | 300 | 300 | 300 | 300 | 300 |
| | N50 (bp) | 805 | 799 | 818 | 734 | 767 | 772 | 709 | 695 | 702 |
| | N90 (bp) | 360 | 360 | 362 | 359 | 365 | 364 | 357 | 356 | 356 |

**Table S3 (xlsx): Relative abundance of the phylotypes at the genus level.** The phylotypes which had relative abundance > 1% in at least one treatment are presented. The designations 0.5, 50, 500, 5,000, 20,000 and 50,000 refer to the $H_2$ mixing ratio in ppmv used for each soil treatment.

**Table S4 (xlsx): Features of all genome bins > 50% completeness and with < 10% contamination.**

**Table S5 (xlsx): Distribution of key metabolic marker genes in 196 MAGs.**

**Table S6 (xlsx): Distribution of metabolic potential of the microbial communities in contaminated soils (Data for Figure 3 and Figure S3).** The designations 0.5, 20,000, and 50,000 denote the different mixing ratios of $H_2$ that each microcosm was treated with (in ppmv).

**Table S7 (xlsx): Distribution of key metabolic marker genes in the 14 MAGs predicted to be capable of PCB degradation.**

**Table S11 (xlsx): Distribution of metabolic genes in the four "high-quality" MAGs (> 90% completeness, < 5% contamination) predicted to be capable of PCB degradation.**

**Figure S1. Copy numbers of (a) whole community 16S rRNA gene and (b) *Dehalobacter* spp. 16S rRNA in the wetland and upland soils.**

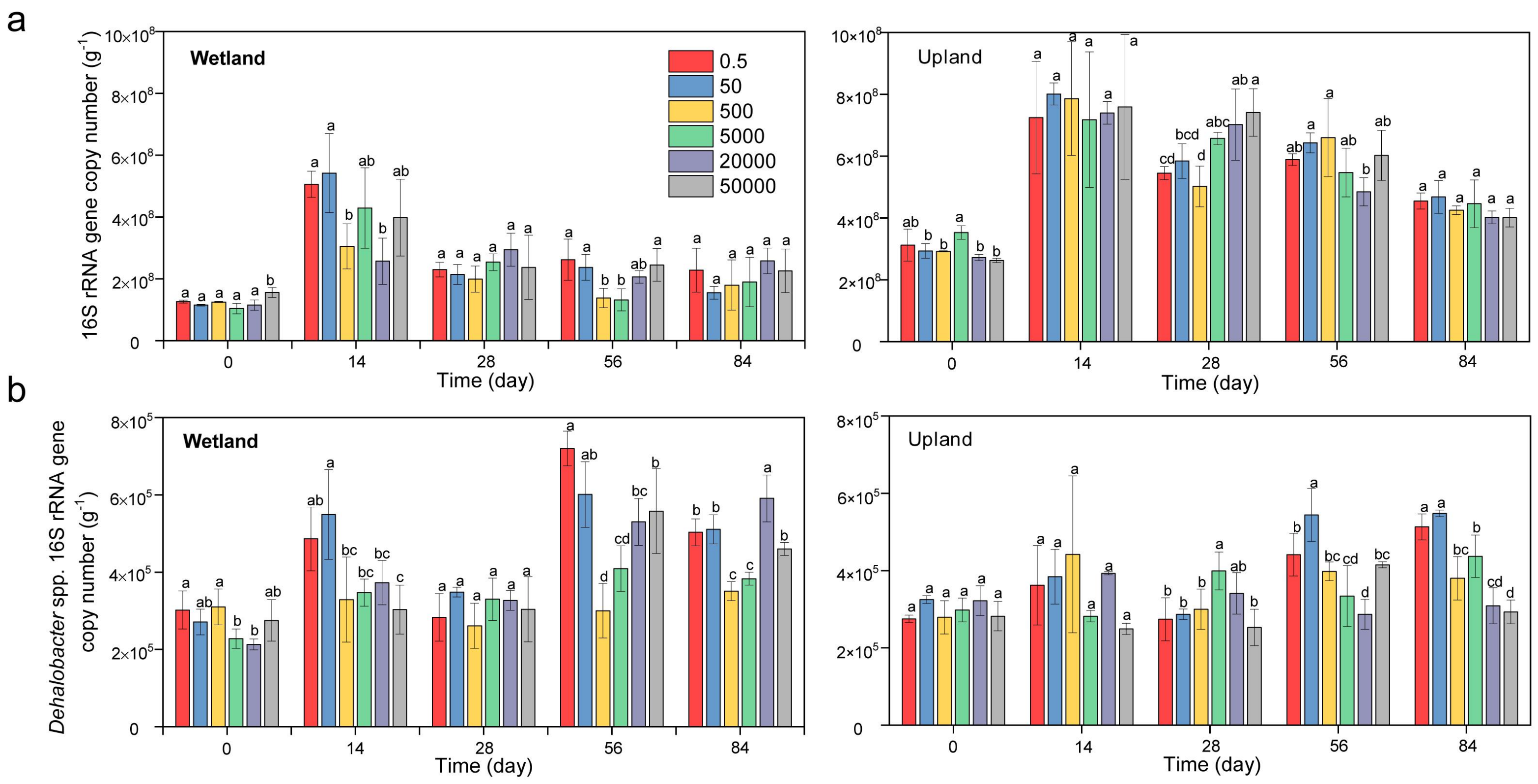

**Figure S2. Box plots comparing Sobs and Chao1 between treatments.** No significant differences were observed between control treatment (0.5 ppmv) and elevated $H_2$ treatment ($p$ > 0.05).

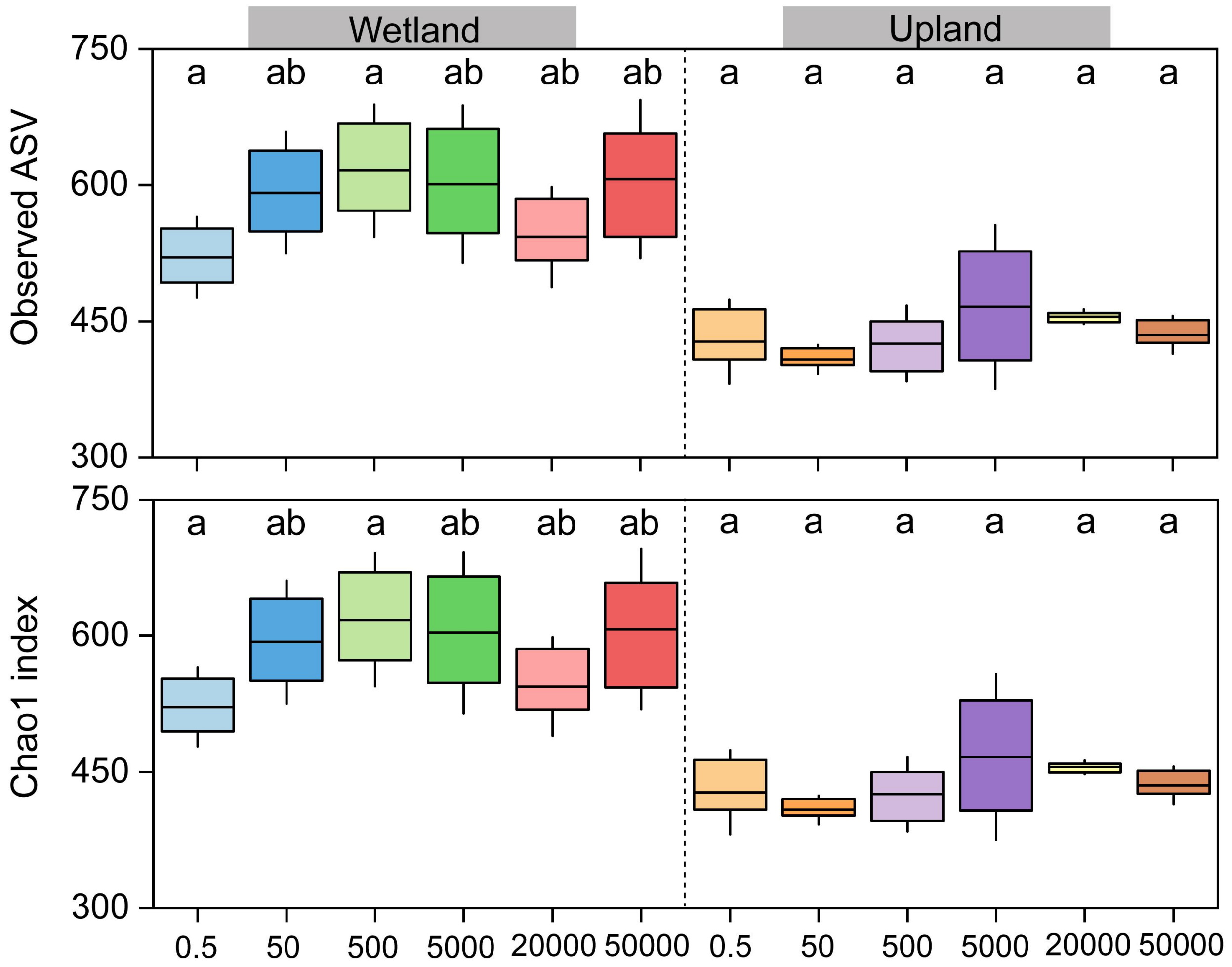

**Figure S3. Ability to utilize inorganic compounds via different pathways in the microbial communities.** Homology-based searches were used to identify signature genes encoding enzymes associated with (from top to bottom): trace gas metabolism, sulfate reduction, nitrogen cycling pathways and other processes. The left heatmap shows the percentage of total community members predicted to encode each signature metabolic gene. To infer abundance, read counts were normalized to gene length and the abundance of single-copy marker genes. The right heatmap shows the presence of these genes across the 196 metagenome-assembled genomes spanning 24 phyla. Abundance was normalized by predicted MAG completeness.

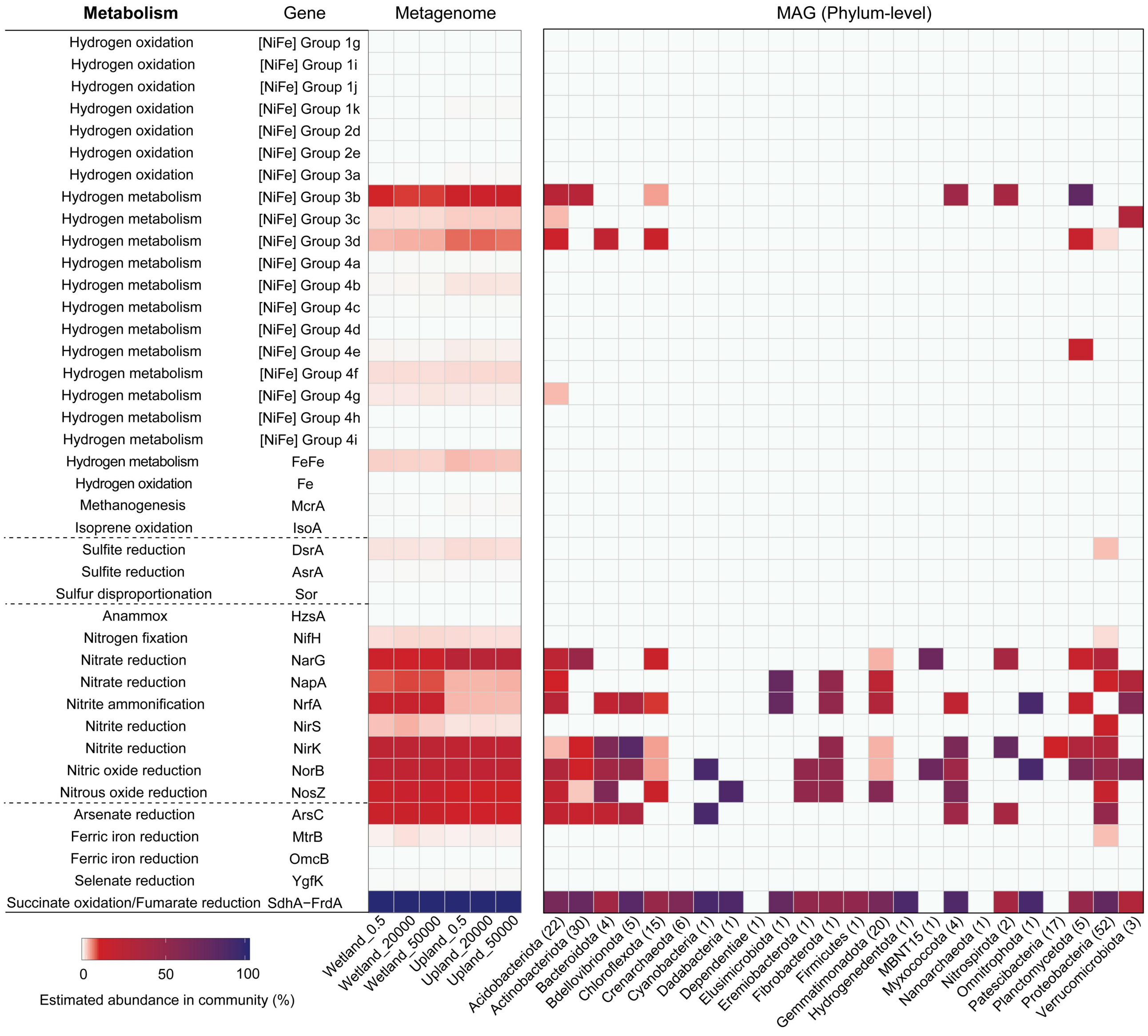
Metabolism
Gene
Metagenome
MAG (Phylum-level)
Hydrogen oxidation [NiFe] Group 1g
Hydrogen oxidation [NiFe] Group 1i
Hydrogen oxidation [NiFe] Group 1j
Hydrogen oxidation [NiFe] Group 1k
Hydrogen oxidation [NiFe] Group 2d
Hydrogen oxidation [NiFe] Group 2e
Hydrogen oxidation [NiFe] Group 3a
Hydrogen metabolism [NiFe] Group 3b
Hydrogen metabolism [NiFe] Group 3c
Hydrogen metabolism [NiFe] Group 3d
Hydrogen metabolism [NiFe] Group 4a
Hydrogen metabolism [NiFe] Group 4b
Hydrogen metabolism [NiFe] Group 4c
Hydrogen metabolism [NiFe] Group 4d
Hydrogen metabolism [NiFe] Group 4e
Hydrogen metabolism [NiFe] Group 4f
Hydrogen metabolism [NiFe] Group 4g
Hydrogen metabolism [NiFe] Group 4h
Hydrogen metabolism [NiFe] Group 4i
Hydrogen metabolism FeFe
Hydrogen oxidation Fe
Methanogenesis McrA
Isoprene oxidation IsoA
Sulfite reduction DsrA
Sulfite reduction AsrA
Sulfur disproportionation Sor
Anammox HzsA
Nitrogen fixation NifH
Nitrate reduction NarG
Nitrate reduction NapA
Nitrite ammonification NrfA
Nitrite reduction NirS
Nitrite reduction NirK
Nitric oxide reduction NorB
Nitrous oxide reduction NosZ
Arsenate reduction ArsC
Ferric iron reduction MtrB
Ferric iron reduction OmcB
Selenate reduction YgfK
Succinate oxidation/Fumarate reduction SdhA−FrdA
Wetland_0.5
Wetland_20000
Wetland_50000
Upland_0.5
Upland_20000
Upland_50000
Acidobacteriota (22)
Actinobacteriota (30)
Bacteroidota (4)
Bdellovibrionota (5)
Chloroflexota (15)
Crenarchaeota (6)
Cyanobacteria (1)
Dadabacteria (1)
Dependentiae (1)
Elusimicrobiota (1)
Eremiobacterota (1)
Fibrobacterota (1)
Firmicutes (1)
Gemmatimonadota (20)
Hydrogenedentota (1)
MBNT15 (1)
Myxococcota (4)
Nanoarchaeota (1)
Nitrospirota (2)
Omnitrophota (1)
Patescibacteria (17)
Planctomycetota (5)
Proteobacteria (52)
Verrucomicrobiota (3)
0
50
100
Estimated abundance in community (%)

**Figure S4.** Maximum-likelihood tree of amino acid sequences of group 1 and 2 [NiFe] hydrogenases, a marker for hydrogen oxidation. The tree shows sequences from metagenome-assembled genomes (blue) from the contaminated soils alongside representative reference sequences (grey). The tree was constructed using the JTT matrix-based model, and was bootstrapped with 50 replicates and midpoint-rooted.

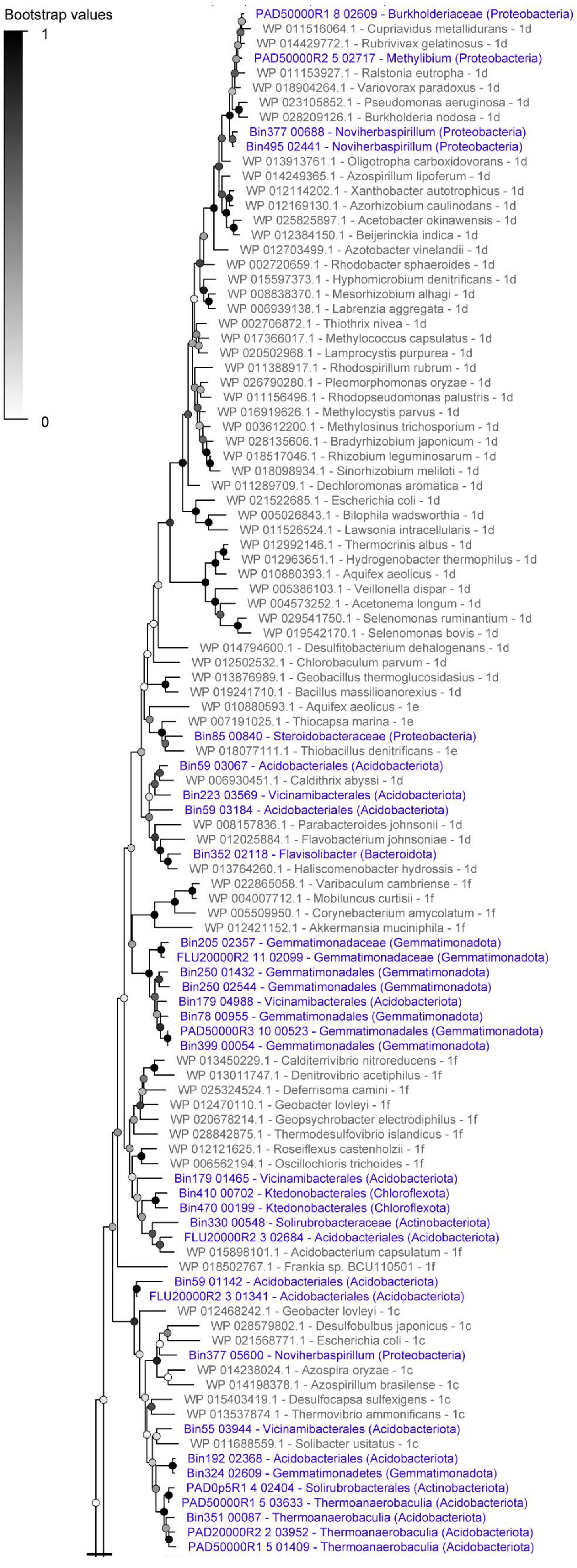
Bootstrap values
1
0
PAD50000R1 8 02609 - Burkholderiaceae (Proteobacteria)
WP 011516064.1 - Cupriavidus metallidurans - 1d
WP 014429772.1 - Rubrivivax gelatinosus - 1d
PAD50000R2 5 02717 - Methylibium (Proteobacteria)
WP 011153927.1 - Ralstonia eutropha - 1d
WP 018904264.1 - Variovorax paradoxus - 1d
WP 023105852.1 - Pseudomonas aeruginosa - 1d
WP 028209126.1 - Burkholderia nodosa - 1d
Bin377 00688 - Noviherbaspirillum (Proteobacteria)
Bin495 02441 - Noviherbaspirillum (Proteobacteria)
WP 013913761.1 - Oligotropha carboxidovorans - 1d
WP 014249365.1 - Azospirillum lipoferum - 1d
WP 012114202.1 - Xanthobacter autotrophicus - 1d
WP 012169130.1 - Azorhizobium caulinodans - 1d
WP 025825897.1 - Acetobacter okinawensis - 1d
WP 012384150.1 - Beijerinckia indica - 1d
WP 012703499.1 - Azotobacter vinelandii - 1d
WP 002720659.1 - Rhodobacter sphaeroides - 1d
WP 015597373.1 - Hyphomicrobium denitrificans - 1d
WP 008838370.1 - Mesorhizobium alhagi - 1d
WP 006939138.1 - Labrenzia aggregata - 1d
WP 002706872.1 - Thiothrix nivea - 1d
WP 017366017.1 - Methylococcus capsulatus - 1d
WP 020502968.1 - Lamprocystis purpurea - 1d
WP 011388917.1 - Rhodospirillum rubrum - 1d
WP 026790280.1 - Pleomorphomonas oryzae - 1d
WP 011156496.1 - Rhodopseudomonas palustris - 1d
WP 016919626.1 - Methylocystis parvus - 1d
WP 003612200.1 - Methylosinus trichosporium - 1d
WP 028135606.1 - Bradyrhizobium japonicum - 1d
WP 018517046.1 - Rhizobium leguminosarum - 1d
WP 018098934.1 - Sinorhizobium meliloti - 1d
WP 011289709.1 - Dechloromonas aromatica - 1d
WP 021522685.1 - Escherichia coli - 1d
WP 005026843.1 - Bilophila wadsworthia - 1d
WP 011526524.1 - Lawsonia intracellularis - 1d
WP 012992146.1 - Thermocrinis albus - 1d
WP 012963651.1 - Hydrogenobacter thermophilus - 1d
WP 010880393.1 - Aquifex aeolicus - 1d
WP 005386103.1 - Veillonella dispar - 1d
WP 004573252.1 - Acetonema longum - 1d
WP 029541750.1 - Selenomonas ruminantium - 1d
WP 019542170.1 - Selenomonas bovis - 1d
WP 014794600.1 - Desulfitobacterium dehalogenans - 1d
WP 012502532.1 - Chlorobaculum parvum - 1d
WP 013876989.1 - Geobacillus thermoglucosidasius - 1d
WP 019241710.1 - Bacillus massilioanorexius - 1d
WP 010880593.1 - Aquifex aeolicus - 1e
WP 007191025.1 - Thiocapsa marina - 1e
Bin85 00840 - Steroidobacteraceae (Proteobacteria)
WP 018077111.1 - Thiobacillus denitrificans - 1e
Bin59 03067 - Acidobacteriales (Acidobacteriota)
WP 006930451.1 - Caldithrix abyssi - 1d
Bin223 03569 - Vicinamibacterales (Acidobacteriota)
Bin59 03184 - Acidobacteriales (Acidobacteriota)
WP 008157836.1 - Parabacteroides johnsonii - 1d
WP 012025884.1 - Flavobacterium johnsoniae - 1d
Bin352 02118 - Flavisolibacter (Bacteroidota)
WP 013764260.1 - Haliscomenobacter hydrossis - 1d
WP 022865058.1 - Varibaculum cambriense - 1f
WP 004007712.1 - Mobiluncus curtisii - 1f
WP 005509950.1 - Corynebacterium amycolatum - 1f
WP 012421152.1 - Akkermansia muciniphila - 1f
Bin205 02357 - Gemmatimonadaceae (Gemmatimonadota)
FLU20000R2 11 02099 - Gemmatimonadaceae (Gemmatimonadota)
Bin250 01432 - Gemmatimonadales (Gemmatimonadota)
Bin250 02544 - Gemmatimonadales (Gemmatimonadota)
Bin179 04988 - Vicinamibacterales (Acidobacteriota)
Bin78 00955 - Gemmatimonadales (Gemmatimonadota)
PAD50000R3 10 00523 - Gemmatimonadales (Gemmatimonadota)
Bin399 00054 - Gemmatimonadales (Gemmatimonadota)
WP 013450229.1 - Calditerrivibrio nitroreducens - 1f
WP 013011747.1 - Denitrovibrio acetiphilus - 1f
WP 025324524.1 - Deferrisoma camini - 1f
WP 012470110.1 - Geobacter lovleyi - 1f
WP 020678214.1 - Geopsychrobacter electrodiphilus - 1f
WP 028842875.1 - Thermodesulfovibrio islandicus - 1f
WP 012121625.1 - Roseiflexus castenholzii - 1f
WP 006562194.1 - Oscillochloris trichoides - 1f
Bin179 01465 - Vicinamibacterales (Acidobacteriota)
Bin410 00702 - Ktedonobacterales (Chloroflexota)
Bin470 00199 - Ktedonobacterales (Chloroflexota)
Bin330 00548 - Solirubrobacteraceae (Actinobacteriota)
FLU20000R2 3 02684 - Acidobacteriales (Acidobacteriota)
WP 015898101.1 - Acidobacterium capsulatum - 1f
WP 018502767.1 - Frankia sp. BCU110501 - 1f
Bin59 01142 - Acidobacteriales (Acidobacteriota)
FLU20000R2 3 01341 - Acidobacteriales (Acidobacteriota)
WP 012468242.1 - Geobacter lovleyi - 1c
WP 028579802.1 - Desulfobulbus japonicus - 1c
WP 021568771.1 - Escherichia coli - 1c
Bin377 05600 - Noviherbaspirillum (Proteobacteria)
WP 014238024.1 - Azospira oryzae - 1c
WP 014198378.1 - Azospirillum brasilense - 1c
WP 015403419.1 - Desulfocapsa sulfexigens - 1c
WP 013537874.1 - Thermovibrio ammonificans - 1c
Bin55 03944 - Vicinamibacterales (Acidobacteriota)
WP 011688559.1 - Solibacter usitatus - 1c
Bin192 02368 - Acidobacteriales (Acidobacteriota)
Bin324 02609 - Gemmatimonadetes (Gemmatimonadota)
PAD0p5R1 4 02404 - Solirubrobacterales (Actinobacteriota)
PAD50000R1 5 03633 - Thermoanaerobaculia (Acidobacteriota)
Bin351 00087 - Thermoanaerobaculia (Acidobacteriota)
PAD20000R2 2 03952 - Thermoanaerobaculia (Acidobacteriota)
PAD50000R1 5 01409 - Thermoanaerobaculia (Acidobacteriota)

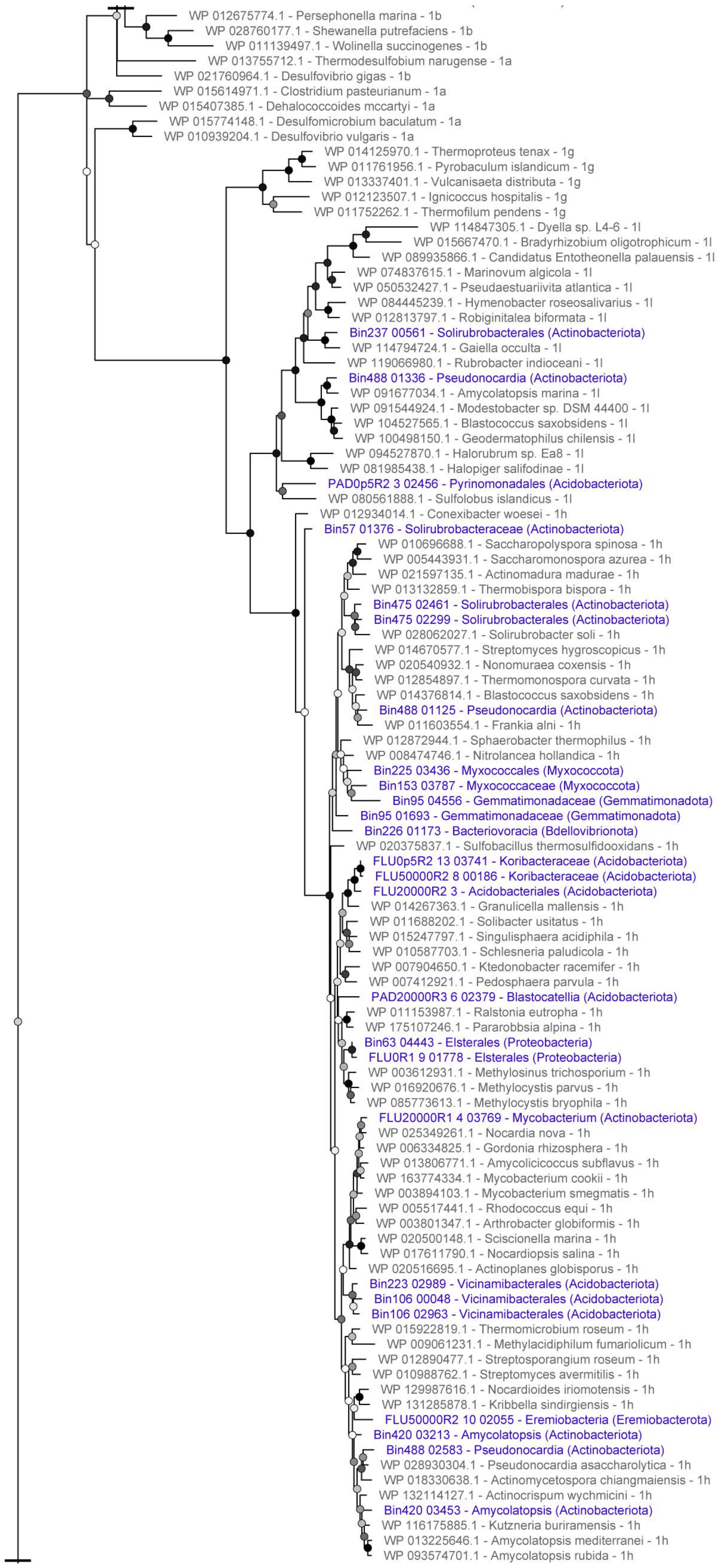

WP 012675774.1 - Persephonella marina - 1b
WP 028760177.1 - Shewanella putrefaciens - 1b
WP 011139497.1 - Wolinella succinogenes - 1b
WP 013755712.1 - Thermodesulfobium narugense - 1a
WP 021760964.1 - Desulfovibrio gigas - 1b
WP 015614971.1 - Clostridium pasteurianum - 1a
WP 015407385.1 - Dehalococcoides mccartyi - 1a
WP 015774148.1 - Desulfomicrobium baculatum - 1a
WP 010939204.1 - Desulfovibrio vulgaris - 1a
WP 014125970.1 - Thermoproteus tenax - 1g
WP 011761956.1 - Pyrobaculum islandicum - 1g
WP 013337401.1 - Vulcanisaeta distributa - 1g
WP 012123507.1 - Ignicoccus hospitalis - 1g
WP 011752262.1 - Thermofilum pendens - 1g
WP 114847305.1 - Dyella sp. L4-6 - 1l
WP 015667470.1 - Bradyrhizobium oligotrophicum - 1l
WP 089935866.1 - Candidatus Entotheonella palauensis - 1l
WP 074837615.1 - Marinovum algicola - 1l
WP 050532427.1 - Pseudaestuariivita atlantica - 1l
WP 084445239.1 - Hymenobacter roseosalivarius - 1l
WP 012813797.1 - Robiginitalea biformata - 1l
Bin237 00561 - Solirubrobacterales (Actinobacteriota)
WP 114794724.1 - Gaiella occulta - 1l
WP 119066980.1 - Rubrobacter indioceani - 1l
Bin488 01336 - Pseudonocardia (Actinobacteriota)
WP 091677034.1 - Amycolatopsis marina - 1l
WP 091544924.1 - Modestobacter sp. DSM 44400 - 1l
WP 104527565.1 - Blastococcus saxobsidens - 1l
WP 100498150.1 - Geodermatophilus chilensis - 1l
WP 094527870.1 - Halorubrum sp. Ea8 - 1l
WP 081985438.1 - Halopiger salifodinae - 1l
PAD0p5R2 3 02456 - Pyrinomonadales (Acidobacteriota)
WP 080561888.1 - Sulfolobus islandicus - 1l
WP 012934014.1 - Conexibacter woesei - 1h
Bin57 01376 - Solirubrobacteraceae (Actinobacteriota)
WP 010696688.1 - Saccharopolyspora spinosa - 1h
WP 005443931.1 - Saccharomonospora azurea - 1h
WP 021597135.1 - Actinomadura madurae - 1h
WP 013132859.1 - Thermobispora bispora - 1h
Bin475 02461 - Solirubrobacterales (Actinobacteriota)
Bin475 02299 - Solirubrobacterales (Actinobacteriota)
WP 028062027.1 - Solirubrobacter soli - 1h
WP 014670577.1 - Streptomyces hygroscopicus - 1h
WP 020540932.1 - Nonomuraea coxensis - 1h
WP 012854897.1 - Thermomonospora curvata - 1h
WP 014376814.1 - Blastococcus saxobsidens - 1h
Bin488 01125 - Pseudonocardia (Actinobacteriota)
WP 011603554.1 - Frankia alni - 1h
WP 012872944.1 - Sphaerobacter thermophilus - 1h
WP 008474746.1 - Nitrolancea hollandica - 1h
Bin225 03436 - Myxococcales (Myxococcota)
Bin153 03787 - Myxococcaceae (Myxococcota)
Bin95 04556 - Gemmatimonadaceae (Gemmatimonadota)
Bin95 01693 - Gemmatimonadaceae (Gemmatimonadota)
Bin226 01173 - Bacteriovoracia (Bdellovibrionota)
WP 020375837.1 - Sulfobacillus thermosulfidooxidans - 1h
FLU0p5R2 13 03741 - Koribacteraceae (Acidobacteriota)
FLU50000R2 8 00186 - Koribacteraceae (Acidobacteriota)
FLU20000R2 3 - Acidobacteriales (Acidobacteriota)
WP 014267363.1 - Granulicella mallensis - 1h
WP 011688202.1 - Solibacter usitatus - 1h
WP 015247797.1 - Singulisphaera acidiphila - 1h
WP 010587703.1 - Schlesneria paludicola - 1h
WP 007904650.1 - Ktedonobacter racemifer - 1h
WP 007412921.1 - Pedosphaera parvula - 1h
PAD20000R3 6 02379 - Blastocatellia (Acidobacteriota)
WP 011153987.1 - Ralstonia eutropha - 1h
WP 175107246.1 - Pararobbsia alpina - 1h
Bin63 04443 - Elsterales (Proteobacteria)
FLU0R1 9 01778 - Elsterales (Proteobacteria)
WP 003612931.1 - Methylosinus trichosporium - 1h
WP 016920676.1 - Methylocystis parvus - 1h
WP 085773613.1 - Methylocystis bryophila - 1h
FLU20000R1 4 03769 - Mycobacterium (Actinobacteriota)
WP 025349261.1 - Nocardia nova - 1h
WP 006334825.1 - Gordonia rhizosphera - 1h
WP 013806771.1 - Amycolicicoccus subflavus - 1h
WP 163774334.1 - Mycobacterium cookii - 1h
WP 003894103.1 - Mycobacterium smegmatis - 1h
WP 005517441.1 - Rhodococcus equi - 1h
WP 003801347.1 - Arthrobacter globiformis - 1h
WP 020500148.1 - Sciscionella marina - 1h
WP 017611790.1 - Nocardiopsis salina - 1h
WP 020516695.1 - Actinoplanes globisporus - 1h
Bin223 02989 - Vicinamibacterales (Acidobacteriota)
Bin106 00048 - Vicinamibacterales (Acidobacteriota)
Bin106 02963 - Vicinamibacterales (Acidobacteriota)
WP 015922819.1 - Thermomicrobium roseum - 1h
WP 009061231.1 - Methylacidiphilum fumariolicum - 1h
WP 012890477.1 - Streptosporangium roseum - 1h
WP 010988762.1 - Streptomyces avermitilis - 1h
WP 129987616.1 - Nocardioides iriomotensis - 1h
WP 131285878.1 - Kribbella sindirgiensis - 1h
FLU50000R2 10 02055 - Eremiobacteria (Eremiobacterota)
Bin420 03213 - Amycolatopsis (Actinobacteriota)
Bin488 02583 - Pseudonocardia (Actinobacteriota)
WP 028930304.1 - Pseudonocardia asaccharolytica - 1h
WP 018330638.1 - Actinomycetospora chiangmaiensis - 1h
WP 132114127.1 - Actinocrispum wychmicini - 1h
Bin420 03453 - Amycolatopsis (Actinobacteriota)
WP 116175885.1 - Kutzneria buriramensis - 1h
WP 013225646.1 - Amycolatopsis mediterranei - 1h
WP 093574701.1 - Amycolatopsis rubida - 1h

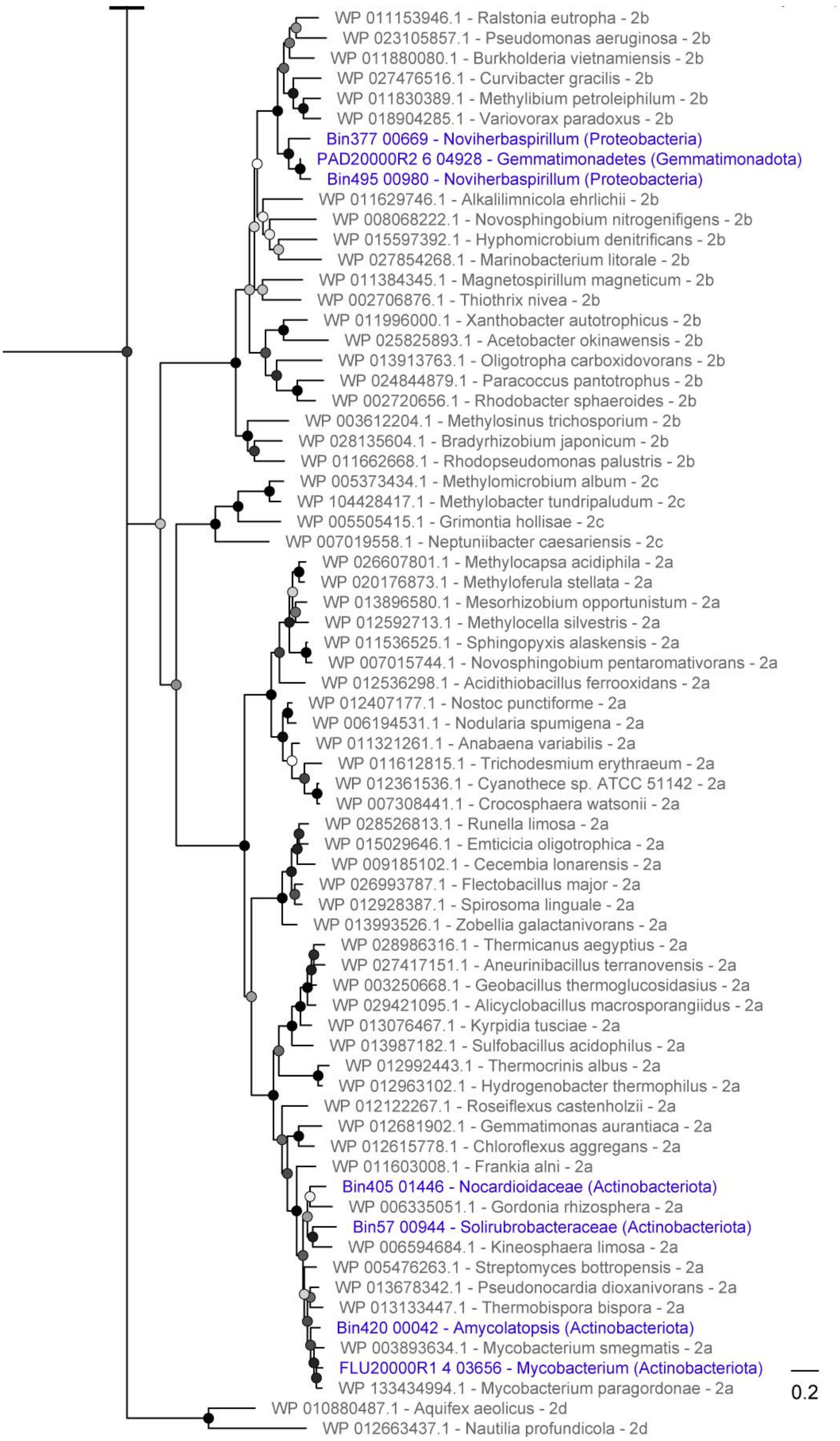

WP 011153946.1 - Ralstonia eutropha - 2b
WP 023105857.1 - Pseudomonas aeruginosa - 2b
WP 011880080.1 - Burkholderia vietnamiensis - 2b
WP 027476516.1 - Curvibacter gracilis - 2b
WP 011830389.1 - Methylibium petroleiphilum - 2b
WP 018904285.1 - Variovorax paradoxus - 2b
Bin377 00669 - Noviherbaspirillum (Proteobacteria)
PAD20000R2 6 04928 - Gemmatimonadetes (Gemmatimonadota)
Bin495 00980 - Noviherbaspirillum (Proteobacteria)
WP 011629746.1 - Alkalilimnicola ehrlichii - 2b
WP 008068222.1 - Novosphingobium nitrogenifigens - 2b
WP 015597392.1 - Hyphomicrobium denitrificans - 2b
WP 027854268.1 - Marinobacterium litorale - 2b
WP 011384345.1 - Magnetospirillum magneticum - 2b
WP 002706876.1 - Thiothrix nivea - 2b
WP 011996000.1 - Xanthobacter autotrophicus - 2b
WP 025825893.1 - Acetobacter okinawensis - 2b
WP 013913763.1 - Oligotropha carboxidovorans - 2b
WP 024844879.1 - Paracoccus pantotrophus - 2b
WP 002720656.1 - Rhodobacter sphaeroides - 2b
WP 003612204.1 - Methylosinus trichosporium - 2b
WP 028135604.1 - Bradyrhizobium japonicum - 2b
WP 011662668.1 - Rhodopseudomonas palustris - 2b
WP 005373434.1 - Methylomicrobium album - 2c
WP 104428417.1 - Methylobacter tundripaludum - 2c
WP 005505415.1 - Grimontia hollisae - 2c
WP 007019558.1 - Neptuniibacter caesariensis - 2c
WP 026607801.1 - Methylocapsa acidiphila - 2a
WP 020176873.1 - Methyloferula stellata - 2a
WP 013896580.1 - Mesorhizobium opportunistum - 2a
WP 012592713.1 - Methylocella silvestris - 2a
WP 011536525.1 - Sphingopyxis alaskensis - 2a
WP 007015744.1 - Novosphingobium pentaromativorans - 2a
WP 012536298.1 - Acidithiobacillus ferrooxidans - 2a
WP 012407177.1 - Nostoc punctiforme - 2a
WP 006194531.1 - Nodularia spumigena - 2a
WP 011321261.1 - Anabaena variabilis - 2a
WP 011612815.1 - Trichodesmium erythraeum - 2a
WP 012361536.1 - Cyanothece sp. ATCC 51142 - 2a
WP 007308441.1 - Crocosphaera watsonii - 2a
WP 028526813.1 - Runella limosa - 2a
WP 015029646.1 - Emticicia oligotrophica - 2a
WP 009185102.1 - Cecembia lonarensis - 2a
WP 026993787.1 - Flectobacillus major - 2a
WP 012928387.1 - Spirosoma linguale - 2a
WP 013993526.1 - Zobellia galactanivorans - 2a
WP 028986316.1 - Thermicanus aegyptius - 2a
WP 027417151.1 - Aneurinibacillus terranovensis - 2a
WP 003250668.1 - Geobacillus thermoglucosidasius - 2a
WP 029421095.1 - Alicyclobacillus macrosporangiidus - 2a
WP 013076467.1 - Kyrpidia tusciae - 2a
WP 013987182.1 - Sulfobacillus acidophilus - 2a
WP 012992443.1 - Thermocrinis albus - 2a
WP 012963102.1 - Hydrogenobacter thermophilus - 2a
WP 012122267.1 - Roseiflexus castenholzii - 2a
WP 012681902.1 - Gemmatimonas aurantiaca - 2a
WP 012615778.1 - Chloroflexus aggregans - 2a
WP 011603008.1 - Frankia alni - 2a
Bin405 01446 - Nocardioidaceae (Actinobacteriota)
WP 006335051.1 - Gordonia rhizosphera - 2a
Bin57 00944 - Solirubrobacteraceae (Actinobacteriota)
WP 006594684.1 - Kineosphaera limosa - 2a
WP 005476263.1 - Streptomyces bottropensis - 2a
WP 013678342.1 - Pseudonocardia dioxanivorans - 2a
WP 013133447.1 - Thermobispora bispora - 2a
Bin420 00042 - Amycolatopsis (Actinobacteriota)
WP 003893634.1 - Mycobacterium smegmatis - 2a
FLU20000R1 4 03656 - Mycobacterium (Actinobacteriota)
WP 133434994.1 - Mycobacterium paragordonae - 2a
0.2
WP 010880487.1 - Aquifex aeolicus - 2d
WP 012663437.1 - Nautilia profundicola - 2d

**Figure S5.** Maximum-likelihood tree of amino acid sequences of ribulose 1,5-bisphosphate carboxylase/oxygenase (RuBisCO) large subunit (RbcL), a marker for carbon fixation through the Calvin-Benson-Bassham (CBB) cycle. The tree shows sequences from metagenome-assembled genomes (blue) from the contaminated soils alongside representative reference sequences (grey). The tree was constructed using the JTT matrix-based model, and was bootstrapped with 50 replicates and midpoint-rooted.

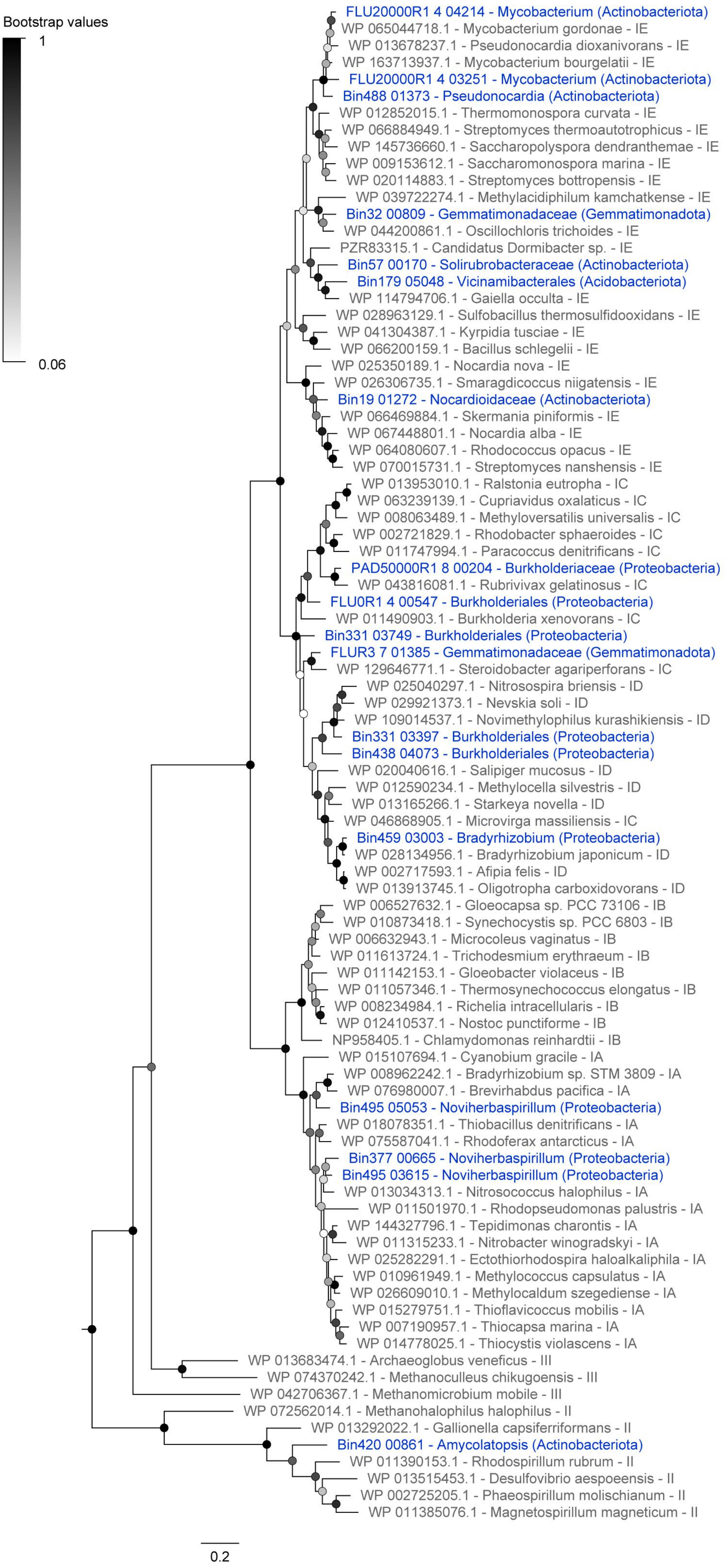

Bootstrap values
1
0.06
FLU20000R1 4 04214 - Mycobacterium (Actinobacteriota)
WP 065044718.1 - Mycobacterium gordonae - IE
WP 013678237.1 - Pseudonocardia dioxanivorans - IE
WP 163713937.1 - Mycobacterium bourgelatii - IE
FLU20000R1 4 03251 - Mycobacterium (Actinobacteriota)
Bin488 01373 - Pseudonocardia (Actinobacteriota)
WP 012852015.1 - Thermomonospora curvata - IE
WP 066884949.1 - Streptomyces thermoautotrophicus - IE
WP 145736660.1 - Saccharopolyspora dendranthemae - IE
WP 009153612.1 - Saccharomonospora marina - IE
WP 020114883.1 - Streptomyces bottropensis - IE
WP 039722274.1 - Methylacidiphilum kamchatkense - IE
Bin32 00809 - Gemmatimonadaceae (Gemmatimonadota)
WP 044200861.1 - Oscillochloris trichoides - IE
PZR83315.1 - Candidatus Dormibacter sp. - IE
Bin57 00170 - Solirubrobacteraceae (Actinobacteriota)
Bin179 05048 - Vicinamibacterales (Acidobacteriota)
WP 114794706.1 - Gaiella occulta - IE
WP 028963129.1 - Sulfobacillus thermosulfidooxidans - IE
WP 041304387.1 - Kyrpidia tusciae - IE
WP 066200159.1 - Bacillus schlegelii - IE
WP 025350189.1 - Nocardia nova - IE
WP 026306735.1 - Smaragdicoccus niigatensis - IE
Bin19 01272 - Nocardioidaceae (Actinobacteriota)
WP 066469884.1 - Skermania piniformis - IE
WP 067448801.1 - Nocardia alba - IE
WP 064080607.1 - Rhodococcus opacus - IE
WP 070015731.1 - Streptomyces nanshensis - IE
WP 013953010.1 - Ralstonia eutropha - IC
WP 063239139.1 - Cupriavidus oxalaticus - IC
WP 008063489.1 - Methyloversatilis universalis - IC
WP 002721829.1 - Rhodobacter sphaeroides - IC
WP 011747994.1 - Paracoccus denitrificans - IC
PAD50000R1 8 00204 - Burkholderiaceae (Proteobacteria)
WP 043816081.1 - Rubrivivax gelatinosus - IC
FLU0R1 4 00547 - Burkholderiales (Proteobacteria)
WP 011490903.1 - Burkholderia xenovorans - IC
Bin331 03749 - Burkholderiales (Proteobacteria)
FLUR3 7 01385 - Gemmatimonadaceae (Gemmatimonadota)
WP 129646771.1 - Steroidobacter agariperforans - IC
WP 025040297.1 - Nitrosospira briensis - ID
WP 029921373.1 - Nevskia soli - ID
WP 109014537.1 - Novimethylophilus kurashikiensis - ID
Bin331 03397 - Burkholderiales (Proteobacteria)
Bin438 04073 - Burkholderiales (Proteobacteria)
WP 020040616.1 - Salipiger mucosus - ID
WP 012590234.1 - Methylocella silvestris - ID
WP 013165266.1 - Starkeya novella - ID
WP 046868905.1 - Microvirga massiliensis - IC
Bin459 03003 - Bradyrhizobium (Proteobacteria)
WP 028134956.1 - Bradyrhizobium japonicum - ID
WP 002717593.1 - Afipia felis - ID
WP 013913745.1 - Oligotropha carboxidovorans - ID
WP 006527632.1 - Gloeocapsa sp. PCC 73106 - IB
WP 010873418.1 - Synechocystis sp. PCC 6803 - IB
WP 006632943.1 - Microcoleus vaginatus - IB
WP 011613724.1 - Trichodesmium erythraeum - IB
WP 011142153.1 - Gloeobacter violaceus - IB
WP 011057346.1 - Thermosynechococcus elongatus - IB
WP 008234984.1 - Richelia intracellularis - IB
WP 012410537.1 - Nostoc punctiforme - IB
NP958405.1 - Chlamydomonas reinhardtii - IB
WP 015107694.1 - Cyanobium gracile - IA
WP 008962242.1 - Bradyrhizobium sp. STM 3809 - IA
WP 076980007.1 - Brevirhabdus pacifica - IA
Bin495 05053 - Noviherbaspirillum (Proteobacteria)
WP 018078351.1 - Thiobacillus denitrificans - IA
WP 075587041.1 - Rhodoferax antarcticus - IA
Bin377 00665 - Noviherbaspirillum (Proteobacteria)
Bin495 03615 - Noviherbaspirillum (Proteobacteria)
WP 013034313.1 - Nitrosococcus halophilus - IA
WP 011501970.1 - Rhodopseudomonas palustris - IA
WP 144327796.1 - Tepidimonas charontis - IA
WP 011315233.1 - Nitrobacter winogradskyi - IA
WP 025282291.1 - Ectothiorhodospira haloalkaliphila - IA
WP 010961949.1 - Methylococcus capsulatus - IA
WP 026609010.1 - Methylocaldum szegediense - IA
WP 015279751.1 - Thioflavicoccus mobilis - IA
WP 007190957.1 - Thiocapsa marina - IA
WP 014778025.1 - Thiocystis violascens - IA
WP 013683474.1 - Archaeoglobus veneficus - III
WP 074370242.1 - Methanoculleus chikugoensis - III
WP 042706367.1 - Methanomicrobium mobile - III
WP 072562014.1 - Methanohalophilus halophilus - II
WP 013292022.1 - Gallionella capsiferriformans - II
Bin420 00861 - Amycolatopsis (Actinobacteriota)
WP 011390153.1 - Rhodospirillum rubrum - II
WP 013515453.1 - Desulfovibrio aespoeensis - II
WP 002725205.1 - Phaeospirillum molischianum - II
WP 011385076.1 - Magnetospirillum magneticum - II
0.2

**Figure S6. Overview of key metabolic capabilities of the MAG predicted to be capable of PCB degradation.** Numbers of genes per genome matching the annotation are showed in the heatmap.

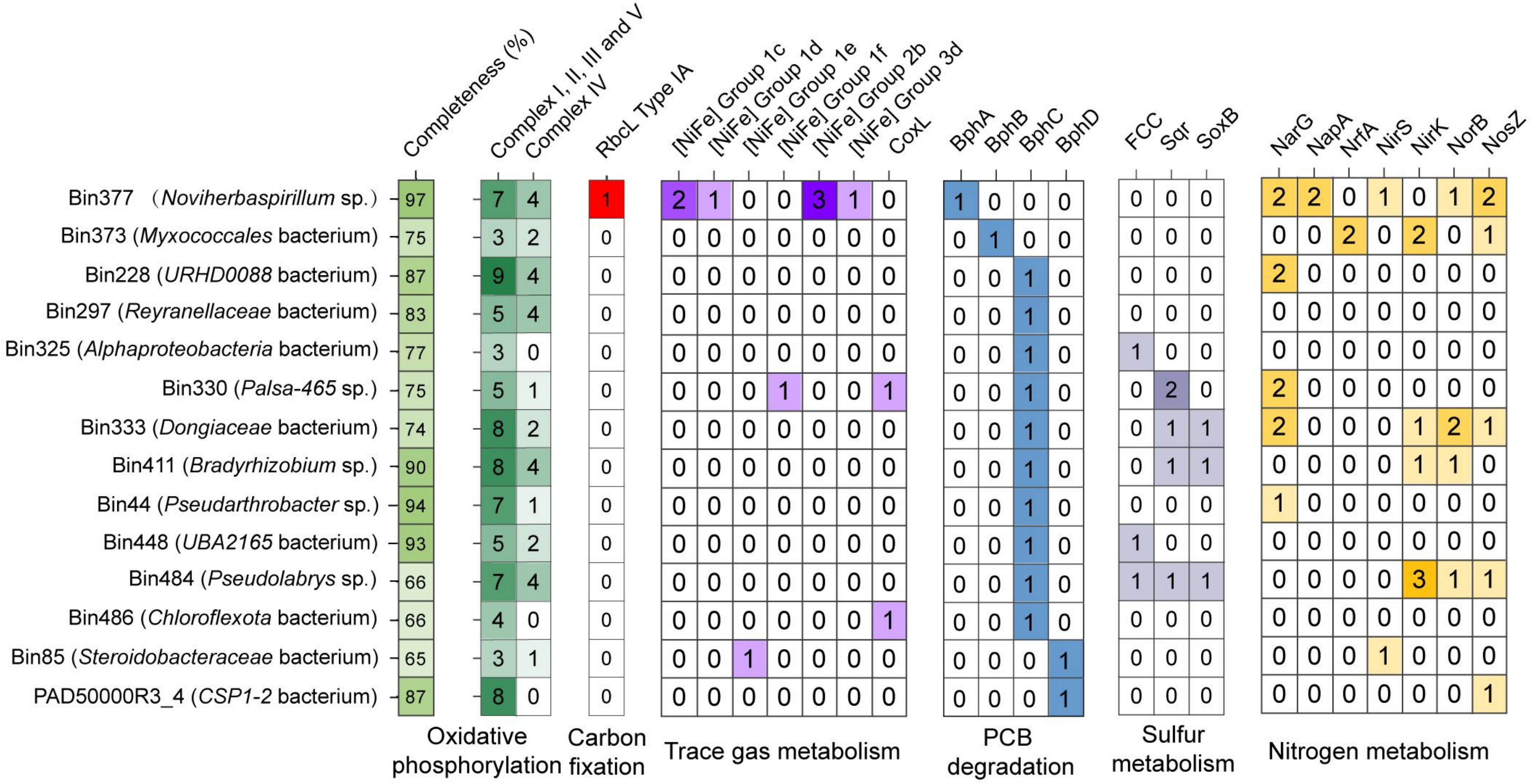

**Figure S7. Residual PCB77 levels in the sterile control microcosms during the incubation period.**

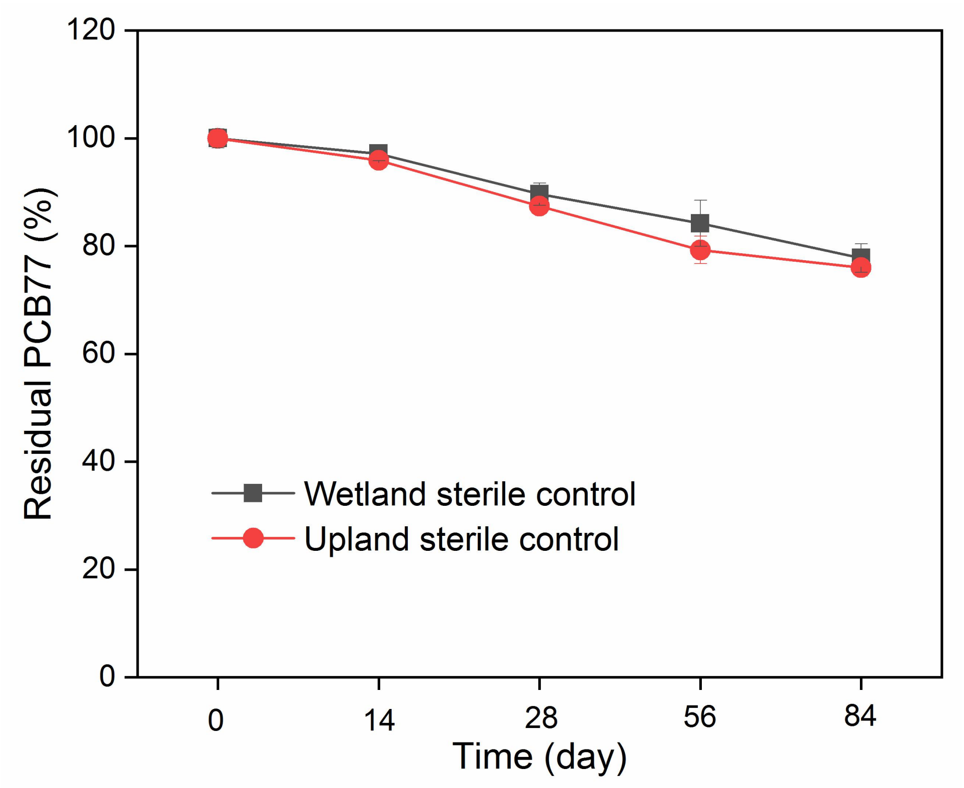
